# A Cofactor-Activated Molecular Switch for Condensate Biogenesis and Catalysis *in Escherichia coli*

**DOI:** 10.64898/2026.06.23.734084

**Authors:** M. Imtiazur Rahman, Roman C. Fabry, Abesh Banerjee, Giovanna Ghirlanda

## Abstract

Biomolecular condensates formed through liquid–liquid phase separation (LLPS) compartmentalize biochemical reactions without enclosing membranes, enabling spatiotemporal control over diverse cellular processes. Engineering genetically encoded proteins that phase separate in response to defined chemical inputs remains a central challenge for synthetic biology. Here, we report a coiled-coil peptide polymer, M1, that undergoes cofactor-dependent condensation both *in vitro* and in *Escherichia coli*. M1 is an ABA triblock construct comprising two terminal helical domains connected by a flexible, intrinsically disordered linker. The terminal domains are derived from a heme-responsive coiled-coil motif that is destabilized in the apo state but assembles into a four-helix bundle upon metalloporphyrin coordination. We demonstrate that M1 forms condensates exclusively in its cofactor-bound state, both *in vitro* and in cells. In *E. coli,* these intracellular condensates accumulate at the cell poles in a concentration-dependent manner. Depletion of cellular heme biosynthetic capacity suppressed condensate formation, which was rescued by supplementation with the heme precursor δ-aminolevulinic acid (δ-ALA) and iron, consistent with metalloporphyrin coordination triggering assembly. The condensates retain peroxidase activity characteristic of heme-containing proteins and catalyze the oxidation of Amplex Red to resorufin both *in vitro* and in living cells. These results establish metalloporphyrin binding as a molecular switch for condensate biogenesis in a structured peptide polymer, directly coupling cofactor coordination, mesoscale assembly, and catalytic function within a single designed system.

**SYNOPSIS:** 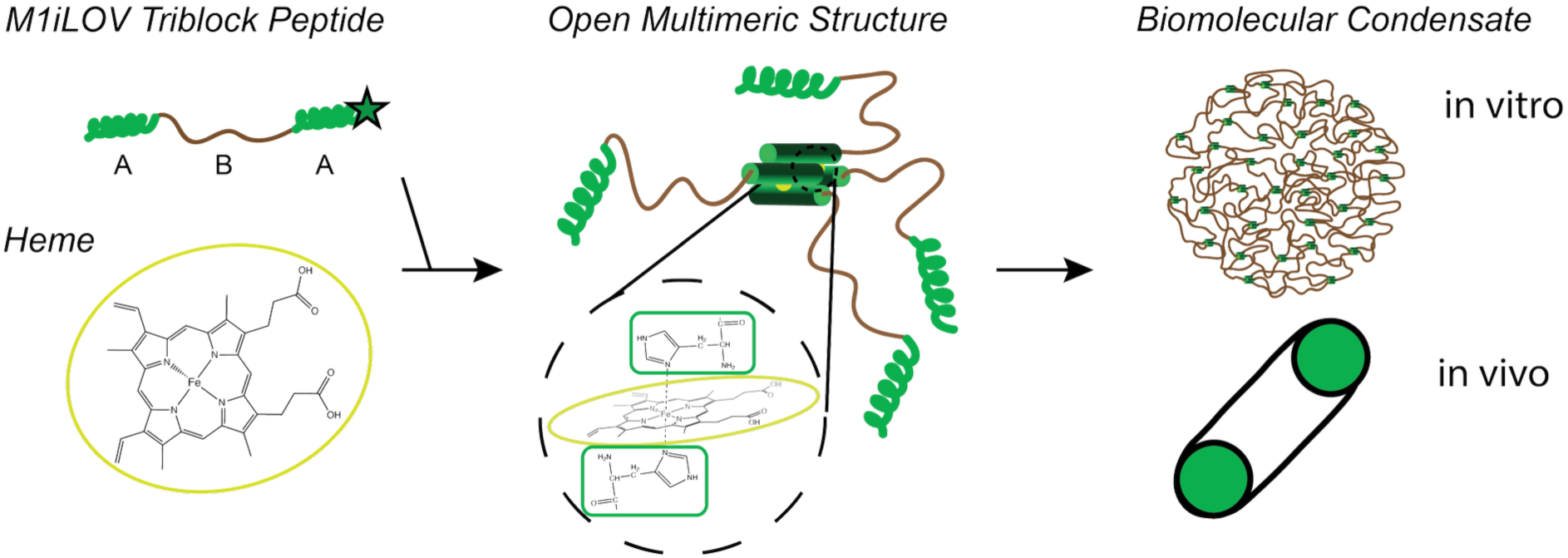

## Introduction

Biomolecular condensates formed through liquid–liquid phase separation dynamically compartmentalize biochemical processes without membrane boundaries and have emerged as a central organizing principle in cell biology.^1–3^ These assemblies concentrate selected biomolecules, buffer compositional fluctuations, and spatially segregate reactions, thereby providing robust spatiotemporal control over cellular functions.^4–7^

The molecular grammar underlying condensate formation is most commonly described by the sticker–spacer framework, in which phase separation is driven by weak, multivalent intermolecular interactions between adhesive domains connected by flexible, disordered sequence regions.^8–14^ Synthetic biology has embraced this principle, constructing engineered condensates from intrinsically disordered polypeptides, elastin-like proteins, and modular sticker–spacer architectures in which multivalent interaction strength and valency are systematically tunable.^15–27^ Structured peptide assemblies represent a comparatively underexplored design space; in principle, these systems allow the geometry, stoichiometry, and intermolecular connectivity of condensate-forming units to be defined with atomic precision, with direct implications for engineering condensates with predictable material properties and switchable assembly behavior.^28–31^Coiled–coil peptides are particularly attractive building blocks for this purpose because their folding, oligomerization state, and intermolecular interactions can be precisely engineered.^32–36^ ABA triblock peptide polymers, in which two coiled-coil sticker domains flank a flexible disordered linker, exploit the defined associative geometry of coiled-coil domains to drive concentration-dependent assembly.^37–42,42–47^ These triblock architectures populate a physicochemical landscape that encompasses soluble oligomers, phase-separated condensates, and percolated hydrogel networks depending on valency, interaction strength, and molecular connectivity.^28–30,48,49^ Most existing coiled-coil triblock systems rely on constitutively associating sticker domains: assembly is concentration-driven but not externally switchable. Coupling coiled-coil folding to a diffusible molecular signal would enable a fundamentally different design principle, in which an exogenous input is transduced directly into mesoscale structural reorganization through stimulus-induced sticker activation.

To address this gap, we developed M1, a genetically encoded ABA triblock polymer in which two identical heme-responsive coiled-coil domains flank a flexible, disordered linker.^40^ Each terminal block is derived from the D2-heme peptide, a designed coiled-coil sequence that is unfolded in the apo state but folds cooperatively into an antiparallel four-helix bundle upon coordination of two heme molecules per bundle.^50^ The central B-domain linker is predicted by IUPred3^51^ to be highly disordered, ensuring conformational flexibility between the terminal associative blocks. Heme-bound M1 self-assembles into hydrogel-like materials above 5 mM *in vitro*, consistent with prior coiled-coil triblock systems.^40^

Here, we demonstrate that cofactor-induced coiled-coil assembly functions as a molecular switch for condensation, enabling M1 to form dynamic condensates both *in vitro* and in *E. coli*. In bacterial cells, M1 forms concentration-dependent polar foci whose assembly depends on heme biosynthesis, whereas a single-domain control construct remains diffused. The resulting condensates retain heme-dependent peroxidase activity, directly coupling cofactor-triggered assembly to catalytic function within an artificial intracellular compartment. These results establish metalloporphyrin binding as a mechanism for activating condensation in structured peptide polymers and provide a generalizable strategy for engineering metal-responsive biomolecular condensates with compartment-localized catalytic activity.

## Results and Discussion

### M1 Forms Dynamic Intracellular Condensates in E. coli

To assess intracellular expression and localization, we fused the fluorescent reporter iLOV to the C terminus of the M1 sequence; as a control, we designed a construct containing a single D2 domain fused to the fluorescent reporter, M1SH-iLOV. M1SH-iLOV can coordinate heme and form a discrete, closed four-helix bundle, while M1 can simultaneously participate in two independent tetrameric nodes and generate an open, percolating network whose formation is the prerequisite for phase separation (Figure 1A). The two constructs were expressed in *E. coli* BL21(DE3) cells under IPTG control. Confocal imaging showed that M1–iLOV formed discrete condensates near the cell poles (Figure 1B). The fraction of M1–iLOV-expressing cells containing detectable foci increased steeply between 2 and 4 hours post-induction, consistent with a progressive accumulation of intracellular protein concentration across the cell population toward and above a critical concentration threshold (Figure S3). In contrast, cells expressing M1SH–iLOV exhibited diffuse fluorescence throughout the cytoplasm and lacked detectable foci, despite comparable overall fluorescence levels (Figure S3). After six hours, over 90% of cells expressing M1-iLOV contained detectable foci (n=216) (Fig. S3); M1-iLOV was enriched 3.2-fold in foci relative to the surrounding cytoplasm (n=274).

**Figure 1.**
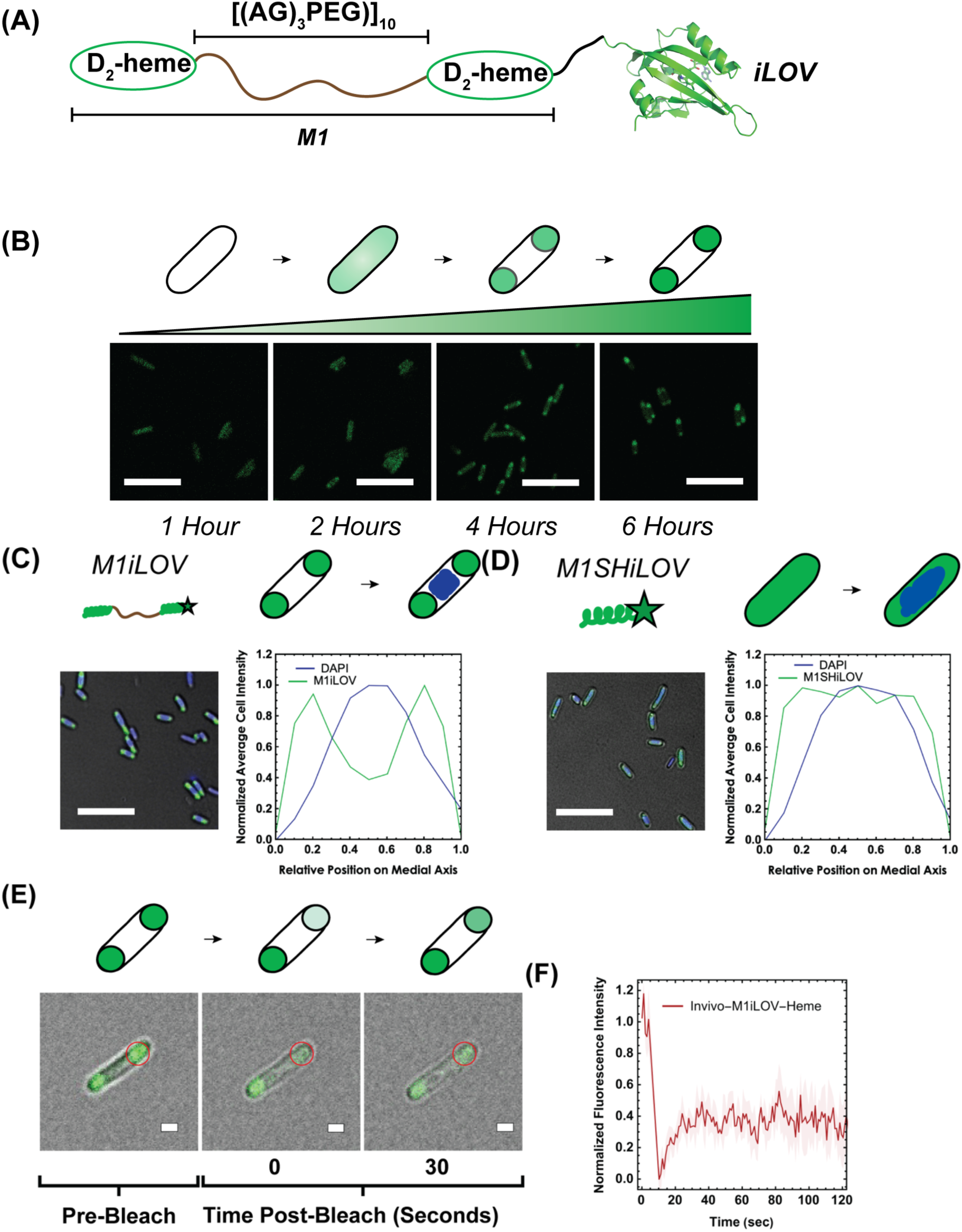
M1-iLOV forms foci in E. coli. (A) M1-iLOV design consists of two conditional helical blocks, D2, linked by ten repeats of a disordered linker (AGAGAGPEG) and tagged by the fluorescent protein iLOV. (B) Merged fluorescence and bright-field microscopy images of *E. coli* cells upon expression of M1-iLOV; scale bars, 5 μm. Cells were grown in LB medium at 37 °C and induced with 500 μM IPTG, and cell samples were taken every 2 h for confocal imaging. (C) Merged fluorescence and bright field microscopy images of cells expressing M1-iLOV and (D) M1SH-iLOV; scale bars, 5 μm. Cells were imaged 4 h post-induction and stained with the DNA dye DAPI (blue). Fluorescence distribution curve of the iLOV channel (green) and of the DAPI channel (blue) plotted vs the medial axis shows nucleoid occlusion for cells expressing M1-iLOV (C, average of 56 cells) and diffuse distribution for cells expressing M1SH-iLOV (D, average of 132 cells). (E) FRAP of M1-iLOV condensates in cells. The red circle indicates the bleached area. (F) Fit to a single exponential yielded t_1/2_ for M1iLOV 4.16 ± 1.41 s. Data are represented as mean ± standard error from n = 5 biologically independent cells. Representative images of pre-bleach, post-bleach frame 1 and the final post-bleach frame are shown. Scale bars, 1 μm.

Given the preferential localization of M1–iLOV condensates at the cell poles, we examined their spatial relationship with the bacterial nucleoid using DAPI staining to visualize chromosomal DNA. Axial fluorescence intensity profiles revealed a clear spatial anti-correlation between M1–iLOV condensates, which were concentrated in DNA-depleted polar regions, whereas the nucleoid was confined to the central portion of the cell body (Figure 1C). M1SH–iLOV, by contrast, displayed diffuse fluorescence with substantially greater overlap with the nucleoid-occupied midcell region (Figure 1D). This spatial partitioning is consistent with a nucleoid occlusion mechanism, in which the compacted chromosomal DNA physically excludes large macromolecular assemblies from the cell center, directing their accumulation toward the DNA-depleted poles. This mode of spatial organization has been previously reported for coiled-coil triblock condensates and other bacterial mesoscale assemblies.^26,28,30,52^

To assess the material properties of the intracellular M1–iLOV assemblies, FRAP was performed on individual polar foci (Figure 1E). Fluorescence recovered with a mean mobile fraction of approximately 40% and a half-time of *t*₁/₂ = 4.16 ± 1.1 s (Figure 1F), demonstrating rapid molecular exchange between the condensed and cytoplasmic phases on experimentally accessible timescales. The timescale of the recovery kinetics is inconsistent with solid-like or gel-like material behavior, and indicates that M1–iLOV molecules remain mobile within the condensate.^24,52^ The incomplete recovery plateau is consistent with depletion of the available cytoplasmic reservoir in the confined intracellular volume, given that a substantial fraction of cellular M1–iLOV partitions into the condensed phase at the expression levels examined.^24,28^

### Condensation is Controlled by Heme Biosynthesis

Because D2 domain assembly depends on metalloporphyrin coordination, we investigated whether intracellular condensation of M1–iLOV is causally linked to heme biosynthesis (Figure 2A). Cells were cultured in M9 minimal medium supplemented with FeCl₃, the heme biosynthetic precursor δ-aminolevulinic acid (δ-ALA), or both, and imaged 5 h post-induction. Under unsupplemented conditions, M1–iLOV fluorescence was predominantly diffuse and fewer than 20% of cells contained detectable condensates (Figure 2B, C). Supplementation with FeCl₃ alone produced only a modest increase in condensate formation. Addition of δ-ALA, by contrast, markedly elevated the condensate-containing fraction to approximately 60%, and combined δ-ALA and FeCl₃ supplementation yielded a comparable result (∼65%), suggesting that porphyrin biosynthetic capacity, rather than iron availability, is the principal limiting factor for condensate assembly under these growth conditions. In support of this interpretation, lysates from cells supplemented with both δ-ALA and FeCl₃ exhibited a Soret absorption band at 414 nm characteristic of bis-histidine-coordinated heme, whereas this feature was absent in lysates from unsupplemented cultures (Figure S5). Together, these results establish that intracellular heme production is the primary determinant of M1 condensate formation, consistent with a model in which metalloporphyrin coordination activates D2 folding and intermolecular assembly to drive phase separation of the ABA triblock polymer in the bacterial cytoplasm.

**Figure 2.**
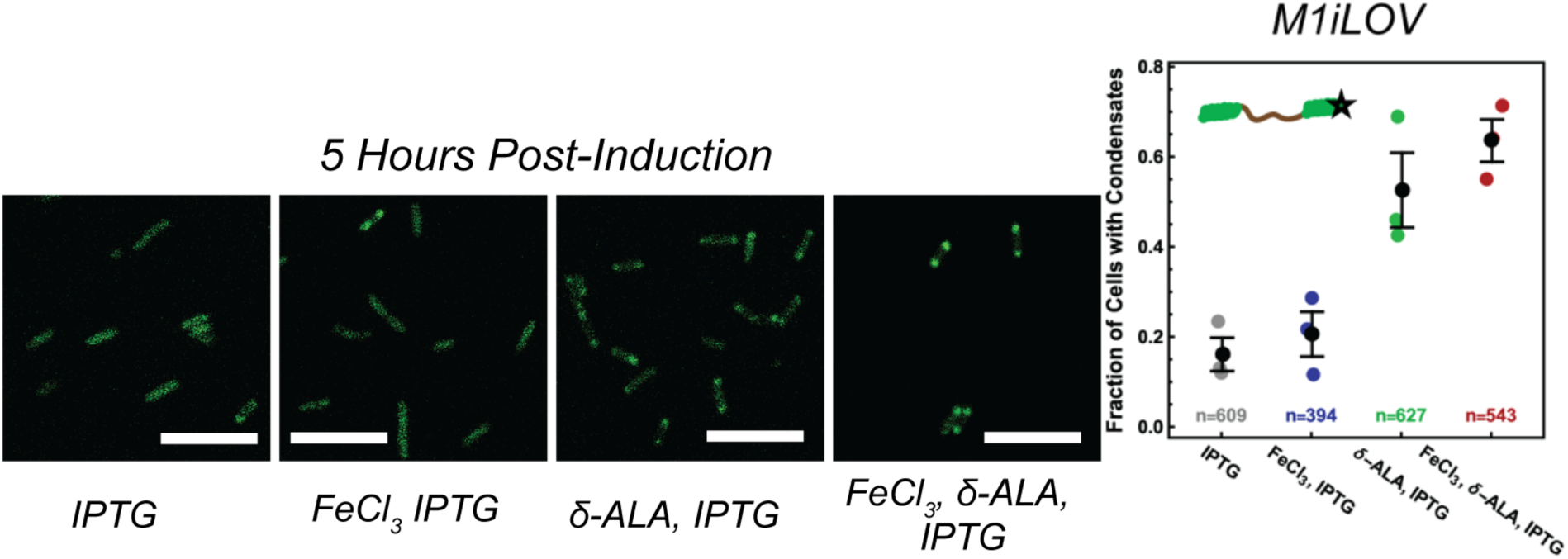
Condensate formation is dependent on heme biosynthesis. (A) Merged fluorescence images of *E.coli* cells expressing M1-iLOV 5 h post-induction with 500 mM IPTG; cells were grown in minimal media (M9) either unsupplemented, or supplemented with (i) ferric chloride, (ii) δ-aminolevulonic acid, or (iii) both ferric chloride and δ-aminolevulonic acid respectively; scale bars, 5 μm. (B) The fraction of cells containing foci is below 20% in unsupplemented conditions (n=609), but increases to 20% with FeCl_3_ (n=394), 55% with δ-ALA (n=627), and 65% with both FeCl_3_ and δ-ALA (n=543).

### Heme binding drives M1-iLOV condensation in vitro

To determine whether condensation is intrinsically driven by cofactor binding rather than by cellular components, we examined M1–iLOV *in vitro* under molecular crowding conditions designed to approximate the bacterial cytoplasm (100 mM sodium glutamate and 9% PEG-8000).^53–56^ In the presence of heme, M1–iLOV formed abundant micron-scale condensates at protein concentrations as low as 100 μM (Figure 3A). In contrast, apo M1–iLOV remained uniformly distributed under identical conditions and did not form detectable condensates. The condensates contained porphyrin, as shown by colocalized fluorescence of the metalloporphyrin and M1–iLOV (Figure 3B), confirming that heme is retained within the condensed phase rather than excluded into the dilute phase.

**Figure 3.**
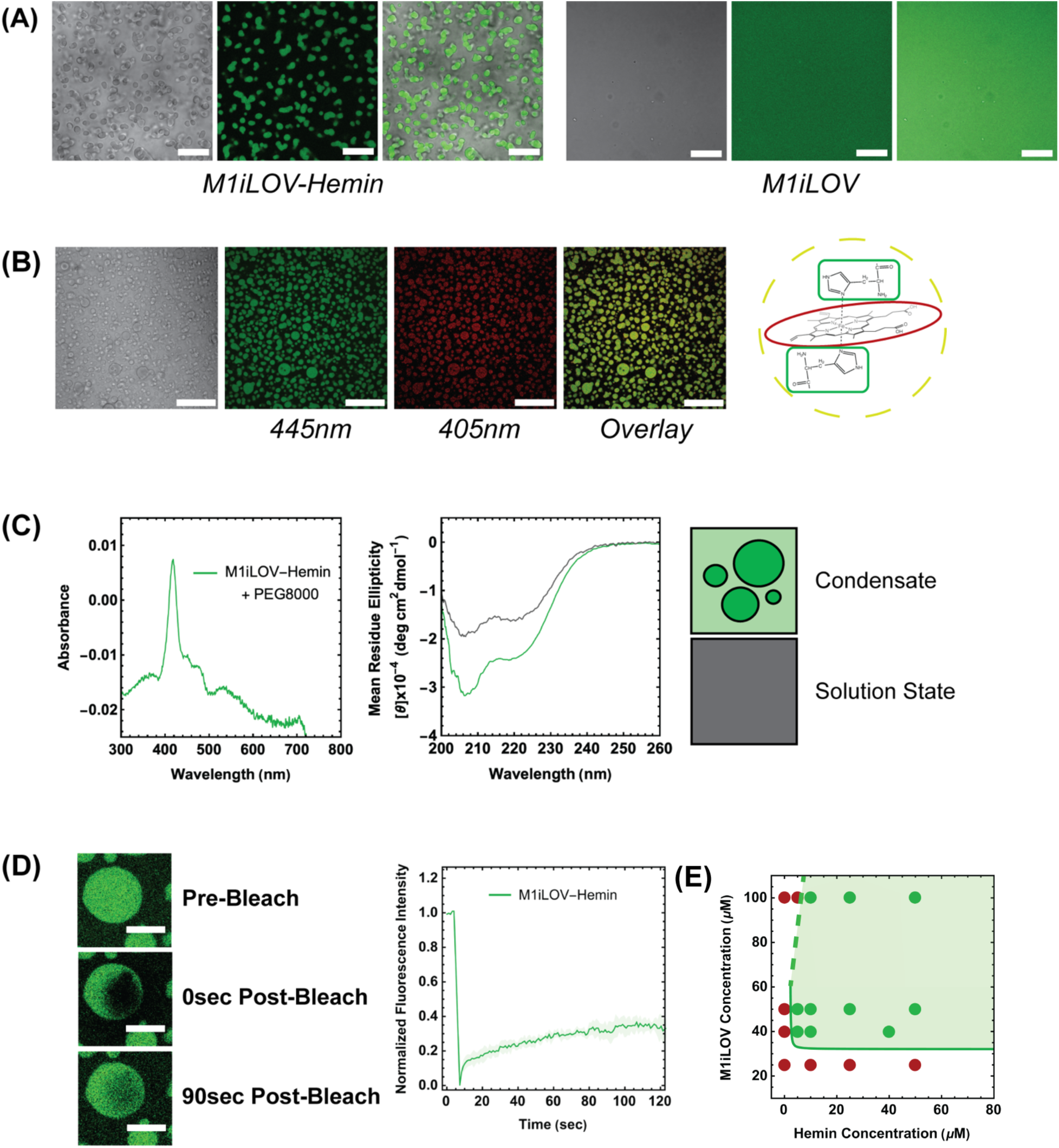
Heme-bound M1-iLOV forms condensates in vitro in crowded solutions. (A) Fluorescent images of M1-iLOV bound to hemin, showing formation of condensates (left) and in the apo state, showing diffuse solution (right); conditions: 100 μM hemin-M1-iLOV, 100 mM sodium glutamate, 9% PEG-8000 in phosphate buffer pH 7.4, scale bars 20 μm. (B) M1-iLOV and ferric isoporphyrin (IHP, a soluble analog of heme) are colocalized in the droplets: from left, brightfield image, iLOV channel, porphyrin channel, merged image; scale bars 50 μm. Conditions: 100 μM M1-iLOV, 350 μM IHP, 100 mM sodium glutamate, 9% PEG-8000 in phosphate buffer pH 7.4. (C) UV-vis spectroscopy of heme-bound M1-iLOV (left) and circular dichroism (right) of apo M1-iLOV in solution (grey line) and heme-bound M1-iLOV in condensates (green line), showing a Soret band max consistent with bis-histidine coordination and increase in helical content upon binding heme in condensates. (D) FRAP of M1-iLOV condensates *in vitro*. Representative images of pre-bleach, post-bleach, and 90 s post-bleach frame (left); scale bars, 10 μm. Data from 30 condensates were analyzed with a double exponential yielding t_1/2_ of 0.449 ± 0.155 s for the fast process and 35.7 ± 7.5 s for the slow process. (E) Phase diagram of M1-iLOV condensation as a function of protein and hemin concentration. Droplets form when the concentration of heme:M1-iLOV complex is sufficiently high. Conditions: 20 mM sodium phosphate, 150mM NaCl, 100 mM sodium glutamate, 9% PEG-8000, pH 7.4.

We next investigated whether the condensed phase contained folded, heme-bound protein. Circular dichroism spectra of apo and hemin-bound M1–iLOV both exhibited minima at 208 and 222 nm characteristic of α-helical secondary structure; because the iLOV domain is itself α-helical, the absolute ellipticity values reflect contributions from both the iLOV fold and the D2 terminal domains (Figure 3C). Hemin binding and condensate formation nonetheless produced a marked increase in ellipticity at 222 nm beyond that attributable to iLOV alone, consistent with cofactor-induced stabilization of the D2 coiled-coil domains upon metalloporphyrin coordination. UV–visible spectroscopy of the condensed fraction revealed the 414 nm Soret band characteristic of bis-histidine-coordinated heme (Figure 3C). FRAP measurements revealed that the *in vitro* condensates remained dynamically exchangeable following assembly (Figure 3D). Fluorescence recovery was biphasic, comprising a rapid intra-droplet component (*t*₁/₂ = 0.45 ± 0.16 s) and a slower inter-phase exchange component (*t*₁/₂ = 35.7 ± 7.5 s), ultimately reaching approximately 40% recovery at 150 s. These recovery kinetics are consistent with liquid-like behavior in which rapid molecular rearrangements within the condensed phase are coupled to slower exchange across the condensate boundary.

Together, these data establish that M1-iLOV condensates are composed of folded, heme-bound protein retaining the native metalloporphyrin coordination geometry previously characterized for the isolated D2 peptide.

To map the concentration requirements for cofactor-driven condensation, we experimentally determined the phase diagram of M1–iLOV as a function of both peptide and hemin concentration. Phase separation was confined to a well-defined two-phase region bounded by a characteristic phase boundary whose shape encodes the dual concentration requirements for assembly (Figure 3E). No condensation was observed at M1–iLOV concentrations of 25 µM across the full range of hemin concentrations tested; droplets were observed at 40 µM, indicating a minimum polymer scaffold density, estimated at 33 µM, below which condensation does not occur regardless of cofactor loading. Above this concentration, condensation depended on hemin: at 100 µM M1–iLOV, phase separation was absent at 5 µM hemin and present at 10 µM hemin, indicating that a minimum cofactor concentration is required to activate sufficient intermolecular connectivity for network formation. Notably, at the lowest hemin concentration tested (5 µM), condensation occurred at intermediate M1–iLOV concentrations (40–50 µM) but not at 100 µM, suggesting that excess apo polymer dilutes the fractional occupancy of heme-loaded sticker domains below the percolation threshold.

This phase behavior is consistent with the sticker–spacer architecture of M1-iLOV, in which two distinct molecular requirements must be simultaneously satisfied. Heme binding must activate a sufficient fraction of terminal D2 sticker domains to generate the intermolecular connectivity required for phase separation; at limiting hemin, most sticker domains remain apo and non-associating, and cross-linking is insufficient regardless of total polymer concentration. Independently, the polymer concentration must exceed c_sat to provide the scaffold density at which activated sticker–sticker contacts generate a network capable of sustaining a macroscopic condensed phase. The resulting phase boundary has a steep vertical arm at low hemin, where cofactor availability limits sticker activation, and a flat horizontal arm at high hemin, where polymer scaffold density becomes the sole determinant of phase separation. Finally, condensate formation under crowding conditions occurred at M1–iLOV concentrations nearly two orders of magnitude below those required for hydrogel formation by M1 in dilute solution,^40^ indicating that molecular crowding strongly lowers the concentration threshold for mesoscale assembly and redirects it toward finite, liquid-like droplets rather than the percolated network characteristic of the hydrogel regime.^55,56^

Collectively, these experiments establish mechanistic coupling between heme binding, coiled-coil folding, and condensation *in vitro*. Under crowded conditions, metalloporphyrin coordination triggers folding and oligomerization of the D2 terminal domains and increases intermolecular connectivity, driving formation of a dynamic condensed phase above the saturation concentration. Heme thus functions as the molecular trigger that couples protein folding to mesoscale assembly, with direct implications for the design of condensates whose formation and dissolution can be externally programmed through ligand availability.

### M1-iLOV condensates retain peroxidase activity

To determine whether M1 condensates retain catalytic function following assembly, we examined hydrogen peroxide-dependent oxidation of Amplex Red to the fluorescent product resorufin, a reaction diagnostic of heme-dependent peroxidase activity (Figure 4A, B).^57^ In *E. coli* expressing M1–iLOV under heme-promoting conditions (δ-ALA and FeCl₃ supplementation), resorufin fluorescence was strongly localized to intracellular condensates and colocalized with the M1–iLOV signal (Figure 4C). Cells grown under unsupplemented conditions, in which condensate formation was substantially reduced, exhibited markedly lower and diffuse resorufin fluorescence. These observations establish that catalytically competent heme is retained within the intracellular condensates and that cofactor-triggered assembly is compatible with enzymatic function in living cells.

**Figure 4.**
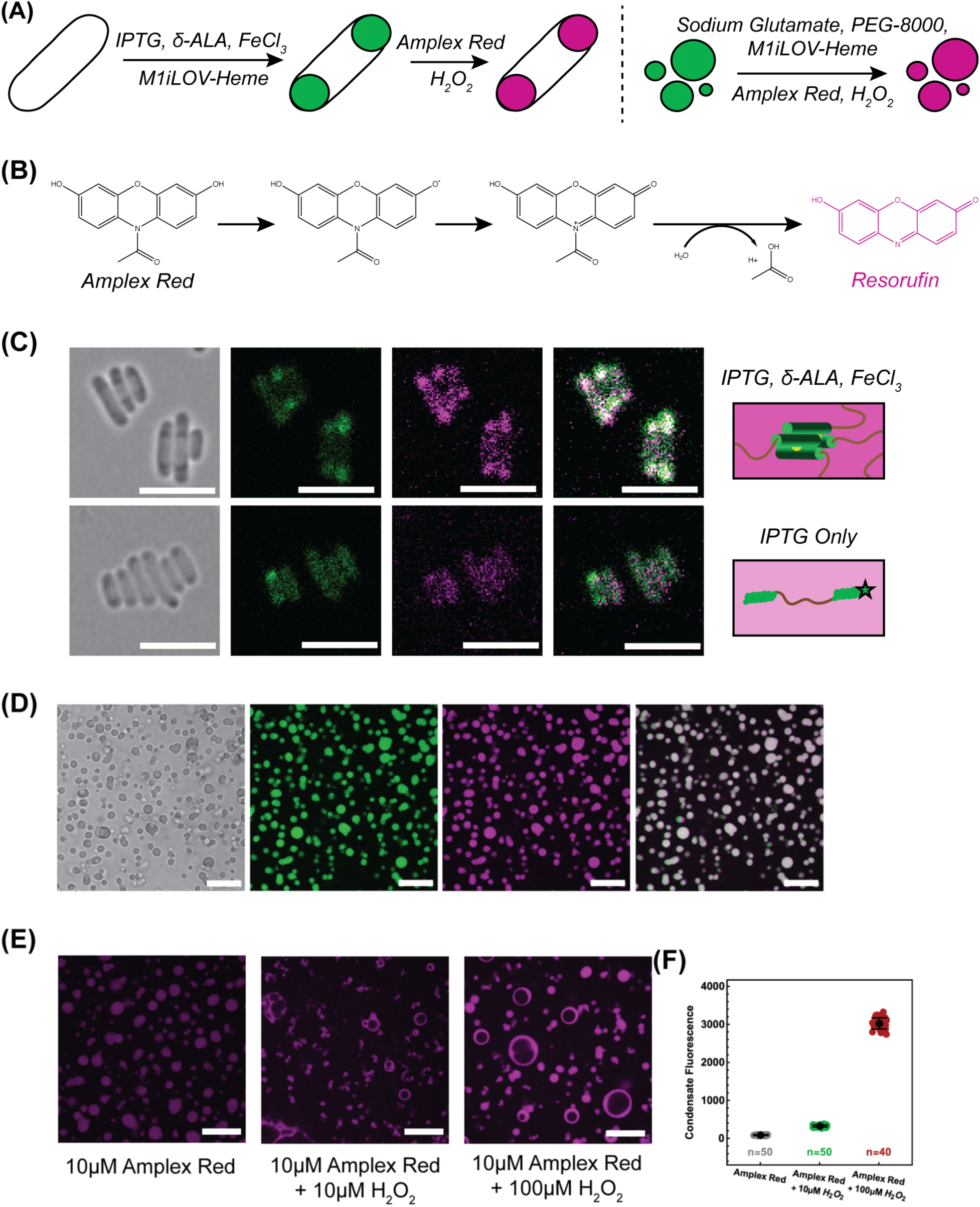
M1-iLOV condensates retain peroxidase activity *in vivo* and *in vitro*. (A) Resorufin formation is associated with the condensates in vivo (left) and in vitro. (B) Simplified progression of Amplex Red substrate as it becomes oxidized to resorufin. Hydrogen peroxide induced radical formation and subsequent substrate oxidation and disproportionation (deacetylation) leading to the formation of a fluorescent resorufin product. (C) Fluorescence images showing heme-containing M1-iLOV condensates retain peroxidase activity in *E. coli.* Resorufin product partitions inside of M1-iLOV condensates showing colocalization of protein and product (top). As a control, M1-iLOV expressed in unsupplemented M9 had much lower resorufin product formation (bottom). (D) Fluorescence images of in vitro heme-containing condensates show similar colocalization of M1-iLOV protein and resorufin product. (E) Resorufin formation in in vitro condensates at increasing hydrogen peroxide concentrations; at higher peroxide concentration, a darker vacuole forms within the droplets. (F) Quantitative analysis of resorufin fluorescence at 0, 10 mM, and 100 mM H_2_O_2_. Conditions: 10 μM Amplex Red, 100 μM M1-iLOV, 100 μM IHP, 100 mM sodium glutamate, 9% PEG-8000 in phosphate buffer pH 7.4.

Catalytic activity was confirmed *in vitro* using condensates formed from IHP-loaded M1–iLOV under crowding conditions. Addition of Amplex Red and hydrogen peroxide produced resorufin fluorescence that colocalized with the condensed phase (Figure 4D), and product formation scaled with H₂O₂ concentration while remaining negligible in its absence (Figure 4E, F), consistent with peroxidase-mediated substrate oxidation. Critically, colocalization of resorufin with the condensate demonstrates that small-molecule substrates access the dense phase and that the elevated local protein density does not sterically occlude the heme centers. At elevated H₂O₂ concentrations, condensates developed internal dark regions not observed prior to catalysis (Figure 4E), suggestive of localized heme degradation or condensate reorganization under sustained oxidative conditions.

These results establish M1 condensates as active biomolecular compartments rather than passive sequestration assemblies. Heme binding serves a dual function within this system: it simultaneously acts as the molecular trigger for condensation and provides the catalytic cofactor required for peroxidase activity, directly coupling the stimulus that drives assembly to the chemical function of the resulting compartment.

## Conclusions

This work establishes cofactor-induced folding as a design principle for stimulus-responsive biomolecular condensate formation in structured peptide polymers. By coupling metalloporphyrin coordination to coiled-coil assembly at the terminal sticker domains of an ABA triblock architecture, M1 undergoes a well-defined folding-to-condensation transition that operates both *in vitro* under physiologically relevant crowding conditions and intracellularly in *E. coli*, where condensate formation is gated by the heme biosynthetic capacity of the cell.

Unlike condensate-forming systems that rely primarily on the multivalency of intrinsically disordered sequences, M1 exploits a cooperative folding transition to generate the intermolecular connectivity required for phase separation. This distinction has material consequences: the condensates are compositionally defined, structurally ordered in the dense phase, and retain catalytically competent heme centers capable of peroxidase activity in both cell-free and cellular contexts. Heme therefore functions in a dual capacity: as the molecular trigger that activates assembly and as the functional cofactor that endows the condensate with enzymatic activity. This dual role directly couples the stimulus driving compartmentalization to the biochemical output of the resulting compartment.

More broadly, these findings establish metalloporphyrins as accessible molecular switches for programming intracellular organization through cofactor-dependent structured assembly, expanding the design vocabulary for synthetic biomolecular condensates beyond intrinsically disordered architectures. The demonstration of ligand-induced folding as a thermodynamic trigger for condensate biogenesis is not specific to heme, and suggests a generalizable strategy for engineering condensates whose formation, dissolution, and activity are regulated by physiologically or therapeutically relevant small molecules, including metal cofactors whose intracellular availability is itself subject to metabolic and homeostatic control.

**Experimental Section. See SI**

## ASSOCIATED CONTENT

### Supporting Information

The Supporting Information is available free of charge on the ACS Publications website. Supplementary methods, figures and tables (file type - PDF))

## AUTHOR INFORMATION

### Author Contributions

MIR, RCF, and AB: investigation, writing (original draft); RCF: data analysis. GG: Conceptualization; investigation; project administration; writing; funding acquisition; supervision. All authors have given approval to the final version of the manuscript.

## ACKNOWLEDGMENT

The authors thank Prof. Tijana Rajh for access to the confocal microscope, and NSF for financial support (NSF EF 1935059).

## ABBREVIATIONS

CD: circular dichroism

## REFERENCES

(1) Banani, S. F.; Lee, H. O.; Hyman, A. A.; Rosen, M. K. Biomolecular Condensates: Organizers of Cellular Biochemistry. Nat Rev Mol Cell Biol 2017, 18 (5), 285–298. 10.1038/nrm.2017.7.

(2) Hyman, A. A.; Weber, C. A.; Jülicher, F. Liquid-Liquid Phase Separation in Biology. Annu. Rev. Cell Dev. Biol. 2014, 30 (1), 39–58. 10.1146/annurev-cellbio-100913-013325.

(3) Boeynaems, S.; Chong, S.; Gsponer, J.; Holt, L.; Milovanovic, D.; Mitrea, D. M.; Mueller-Cajar, O.; Portz, B.; Reilly, J. F.; Reinkemeier, C. D.; Sabari, B. R.; Sanulli, S.; Shorter, J.; Sontag, E.; Strader, L.; Stachowiak, J.; Weber, S. C.; White, M.; Zhang, H.; Zweckstetter, M.; Elbaum-Garfinkle, S.; Kriwacki, R. Phase Separation in Biology and Disease; Current Perspectives and Open Questions. Journal of Molecular Biology 2023, 435 (5), 167971. 10.1016/j.jmb.2023.167971.

(4) Jiang, H.; Todd Stukenberg, P. Emerging Roles of Liquid–Liquid Phase Separation and Membraneless Organelles in Cancer Progression. In Droplets of Life; Elsevier, 2023; pp 651–662. 10.1016/B978-0-12-823967-4.00004-X.

(5) Riggs, C. L.; Ivanov, P. Stress, Membraneless Organelles, and Liquid–Liquid Phase Separation. In Droplets of Life; Elsevier, 2023; pp 505–529. 10.1016/B978-0-12-823967-4.00026-9.

(6) Haider, R.; Boyko, S.; Surewicz, W. K. Liquid–Liquid Phase Separation in Neurodegenerative Diseases. In Droplets of Life; Elsevier, 2023; pp 619–650. 10.1016/B978-0-12-823967-4.00018-X.

(7) Ganser, L. R.; Djaja, N. A.; Myong, S. Biochemical and Structural Biology Aspects of Liquid–Liquid Phase Separation: An Interplay between Proteins and RNA. In Droplets of Life; Elsevier, 2023; pp 133–155. 10.1016/B978-0-12-823967-4.00014-2.

(8) Harmon, T. S.; Holehouse, A. S.; Rosen, M. K.; Pappu, R. V. Intrinsically Disordered Linkers Determine the Interplay between Phase Separation and Gelation in Multivalent Proteins. eLife 2017, 6, e30294. 10.7554/eLife.30294.

(9) Pappu, R. V.; Cohen, S. R.; Dar, F.; Farag, M.; Kar, M. Phase Transitions of Associative Biomacromolecules. Chem. Rev. 2023, 123 (14), 8945–8987. 10.1021/acs.chemrev.2c00814.

(10) Posey, A. E.; Holehouse, A. S.; Pappu, R. V. Phase Separation of Intrinsically Disordered Proteins. Methods in Enzymology 2018, 611, 1–30.

(11) Choi, J.-M.; Holehouse, A. S.; Pappu, R. V. Physical Principles Underlying the Complex Biology of Intracellular Phase Transitions. Annu Rev Biophys 2020, 49, 107–133. 10.1146/annurev-biophys-121219-081629.

(12) Cohan, M. C.; Pappu, R. V. Making the Case for Disordered Proteins and Biomolecular Condensates in Bacteria. Trends in Biochemical Sciences 2020, 45 (8), 668–680. 10.1016/j.tibs.2020.04.011.

(13) Schuster, B. S.; Dignon, G. L.; Tang, W. S.; Kelley, F. M.; Ranganath, A. K.; Jahnke, C. N.; Simpkins, A. G.; Regy, R. M.; Hammer, D. A.; Good, M. C.; Mittal, J. Identifying Sequence Perturbations to an Intrinsically Disordered Protein That Determine Its Phase-Separation Behavior. Proc. Natl. Acad. Sci. U.S.A. 2020, 117 (21), 11421–11431. 10.1073/pnas.2000223117.

(14) Welles, R. M.; Sojitra, K. A.; Garabedian, M. V.; Xia, B.; Wang, W.; Guan, M.; Regy, R. M.; Gallagher, E. R.; Hammer, D. A.; Mittal, J.; Good, M. C. Determinants That Enable Disordered Protein Assembly into Discrete Condensed Phases. Nat. Chem. 2024, 1–11. 10.1038/s41557-023-01423-7.

(15) Simon, J. R.; Carroll, N. J.; Rubinstein, M.; Chilkoti, A.; López, G. P. Programming Molecular Self-Assembly of Intrinsically Disordered Proteins Containing Sequences of Low Complexity. Nature Chem 2017, 9 (6), 509–515. 10.1038/nchem.2715.

(16) Dai, Y.; Farag, M.; Lee, D.; Zeng, X.; Kim, K.; Son, H.; Guo, X.; Su, J.; Peterson, N.; Mohammed, J.; Ney, M.; Shapiro, D. M.; Pappu, R. V.; Chilkoti, A.; You, L. Programmable Synthetic Biomolecular Condensates for Cellular Control. Nat Chem Biol 2023, 19 (4), 518–528. 10.1038/s41589-022-01252-8.

(17) Dai, Y.; You, L.; Chilkoti, A. Engineering Synthetic Biomolecular Condensates. Nat Rev Bioeng 2023, 1 (7), 466–480. 10.1038/s44222-023-00052-6.

(18) Yu, W.; Jin, K.; Wang, D.; Wang, N.; Li, Y.; Liu, Y.; Li, J.; Du, G.; Lv, X.; Chen, J.; Ledesma-Amaro, R.; Liu, L. De Novo Engineering of Programmable and Multi-Functional Biomolecular Condensates for Controlled Biosynthesis. Nat Commun 2024, 15 (1), 7989. 10.1038/s41467-024-52411-5.

(19) Hammer, S. K.; Avalos, J. L. Harnessing Yeast Organelles for Metabolic Engineering. Nat Chem Biol 2017, 13 (8), 823–832. 10.1038/nchembio.2429.

(20) Garabedian, M. V.; Su, Z.; Dabdoub, J.; Tong, M.; Deiters, A.; Hammer, D. A.; Good, M. C. Protein Condensate Formation via Controlled Multimerization of Intrinsically Disordered Sequences. Biochemistry 2022, 61 (22), 2470–2481. 10.1021/acs.biochem.2c00250.

(21) Caldwell, R. M.; Bermudez, J. G.; Thai, D.; Aonbangkhen, C.; Schuster, B. S.; Courtney, T.; Deiters, A.; Hammer, D. A.; Chenoweth, D. M.; Good, M. C. Optochemical Control of Protein Localization and Activity within Cell-like Compartments. Biochemistry 2018, 57 (18), 2590–2596. 10.1021/acs.biochem.8b00131.

(22) Schuster, B. S.; Reed, E. H.; Parthasarathy, R.; Jahnke, C. N.; Caldwell, R. M.; Bermudez, J. G.; Ramage, H.; Good, M. C.; Hammer, D. A. Controllable Protein Phase Separation and Modular Recruitment to Form Responsive Membraneless Organelles. Nat Commun 2018, 9 (1), 2985. 10.1038/s41467-018-05403-1.

(23) Reed, E. H.; Schuster, B. S.; Good, M. C.; Hammer, D. A. SPLIT: Stable Protein Coacervation Using a Light Induced Transition. ACS Synth. Biol. 2020, 9 (3), 500–507. 10.1021/acssynbio.9b00503.

(24) Yeong, V.; Werth, E. G.; Brown, L. M.; Obermeyer, A. C. Formation of Biomolecular Condensates in Bacteria by Tuning Protein Electrostatics. ACS Cent. Sci. 2020, 6 (12), 2301–2310. 10.1021/acscentsci.0c01146.

(25) Horn, J. M.; Zhu, Y.; Ahn, S. Y.; Obermeyer, A. C. Self-Assembly of Globular Proteins with Intrinsically Disordered Protein Polyelectrolytes and Block Copolymers. Soft Matter 2022, 18 (31), 5759–5769. 10.1039/D2SM00415A.

(26) Zu, H.; Pan, F.; Fan, H.-L.; Qian, Z.-G.; Xia, X.-X. Dynamic Structural Organization of Biomolecular Condensates in *Escherichia Coli* Cells. JACS Au 2025, 5 (10), 4728–4739. 10.1021/jacsau.5c00600.

(27) Costantino, M.; Young, E. J.; Banerjee, A.; Kerfeld, C. A.; Ghirlanda, G. Interfacing Bacterial Microcompartment Shell Proteins with Genetically Encoded Condensates. Protein Science 2025, 34 (3), e70061. 10.1002/pro.70061.

(28) Hilditch, A. T.; Romanyuk, A.; Cross, S. J.; Obexer, R.; McManus, J. J.; Woolfson, D. N. Assembling Membraneless Organelles from de Novo Designed Proteins. Nat. Chem. 2024, 16 (1), 89–97. 10.1038/s41557-023-01321-y.

(29) Hilditch, A. T.; Romanyuk, A.; Hodgson, L. R.; Mantell, J.; Neal, C. R.; Verkade, P.; Obexer, R.; Serpell, L. C.; McManus, J. J.; Woolfson, D. N. Maturation and Conformational Switching of a De Novo Designed Phase-Separating Polypeptide. J. Am. Chem. Soc. 2024. 10.1021/jacs.4c00256.

(30) Tomares, D. T.; Mann, M.; DiBernardo, E.; Childers, W. S. Repurposing Peptide Nanomaterials as Synthetic Biomolecular Condensates in Bacteria. ACS Synth. Biol. 2022, 11 (6), 2154–2162. 10.1021/acssynbio.2c00078.

(31) He, Z.; Sommer, J.-U.; Harmon, T. S. Impact of Coiled-Coil Domains on the Phase Behavior of Biomolecular Condensates. ACS Macro Lett. 2025, 14 (4), 413–419. 10.1021/acsmacrolett.4c00821.

(32) Ramšak, M.; Ramirez, D. A.; Hough, L. E.; Shirts, M. R.; Vidmar, S.; Eleršič Filipič, K.; Anderluh, G.; Jerala, R. Programmable de Novo Designed Coiled Coil-Mediated Phase Separation in Mammalian Cells. Nat Commun 2023, 14 (1), 7973. 10.1038/s41467-023-43742-w.

(33) Ramirez, D. A.; Hough, L. E.; Shirts, M. R. Coiled-Coil Domains Are Sufficient to Drive Liquid-Liquid Phase Separation in Protein Models. Biophysical Journal 2024, 123 (6), 703–717. 10.1016/j.bpj.2024.02.007.

(34) Lee, M. J.; Mantell, J.; Hodgson, L.; Alibhai, D.; Fletcher, J. M.; Brown, I. R.; Frank, S.; Xue, W.-F.; Verkade, P.; Woolfson, D. N.; Warren, M. J. Engineered Synthetic Scaffolds for Organizing Proteins within the Bacterial Cytoplasm. Nat Chem Biol 2018, 14 (2), 142–147. 10.1038/nchembio.2535.

(35) Lupas, A. N.; Gruber, M. THE STRUCTURE OF A-HELICAL COILED COILS.

(36) Huang, P.-S.; Boyken, S. E.; Baker, D. The Coming of Age of de Novo Protein Design. Nature 2016, 537 (7620), 320–327. 10.1038/nature19946.

(37) Banwell, E. F.; Abelardo, E. S.; Adams, D. J.; Birchall, M. A.; Corrigan, A.; Donald, A. M.; Kirkland, M.; Serpell, L. C.; Butler, M. F.; Woolfson, D. N. Rational Design and Application of Responsive α-Helical Peptide Hydrogels. Nature Mater 2009, 8 (7), 596–600. 10.1038/nmat2479.

(38) D’Souza, A.; Marshall, L. R.; Yoon, J.; Kulesha, A.; Edirisinghe, D. I. U.; Chandrasekaran, S.; Rathee, P.; Prabhakar, R.; Makhlynets, O. V. Peptide Hydrogel with Self-Healing and Redox-Responsive Properties. Nano Convergence 2022, 9 (1), 18. 10.1186/s40580-022-00309-7.

(39) Dexter, A. F.; Fletcher, N. L.; Creasey, R. G.; Filardo, F.; Boehm, M. W.; Jack, K. S. Fabrication and Characterization of Hydrogels Formed from Designer Coiled-Coil Fibril-Forming Peptides. RSC Adv. 2017, 7 (44), 27260–27271. 10.1039/C7RA02811C.

(40) Halloran, N. R.; Banerjee, A.; Ghirlanda, G. Self-Assembling Peptide–Co-PPIX Complex Catalyzes Photocatalytic Hydrogen Evolution and Forms Hydrogels. Molecules 2025, 30 (8), 1707. 10.3390/molecules30081707.

(41) Meleties, M.; Katyal, P.; Lin, B.; Britton, D.; Kim Montclare, J. Self-Assembly of Stimuli-Responsive Coiled-Coil Fibrous Hydrogels. Soft Matter 2021, 17 (26), 6470–6476. 10.1039/D1SM00780G.

(42) Mout, R.; Bretherton, R. C.; Decarreau, J.; Lee, S.; Gregorio, N.; Edman, N. I.; Ahlrichs, M.; Hsia, Y.; Sahtoe, D. D.; Ueda, G.; Sharma, A.; Schulman, R.; DeForest, C. A.; Baker, D. De Novo Design of Modular Protein Hydrogels with Programmable Intra-and Extracellular Viscoelasticity. Proceedings of the National Academy of Sciences 2024, 121 (6), e2309457121. 10.1073/pnas.2309457121.

(43) Petka, D. A., W. A., Harden, J. L., McGrath, K. P., Wirtz, D. &. Tirrell. Reversible Hydrogels from Self-Assembling Artificial Proteins. Science 1998, 281.

(44) Wang, Q. et al. A Supramolecular-Hydrogel-Encapsulated Hemin as an Artificial Enzyme to Mimic Peroxidase. Angewandte Chemie - International Edition 2007, 46.

(45) Wheeldon, I. R.; Gallaway, J. W.; Barton, S. C.; Banta, S. Bioelectrocatalytic Hydrogels from Electron-Conducting Metallopolypeptides Coassembled with Bifunctional Enzymatic Building Blocks. Proceedings of the National Academy of Sciences 2008, 105 (40), 15275–15280. 10.1073/pnas.0805249105.

(46) Xu, C.; Breedveld, V.; Kopeček, J. Reversible Hydrogels from Self-Assembling Genetically Engineered Protein Block Copolymers. Biomacromolecules 2005, 6 (3), 1739–1749. 10.1021/bm050017f.

(47) Katyal, P.; Mahmoudinobar, F.; Montclare, J. K. Recent Trends in Peptide and Protein-Based Hydrogels. Current Opinion in Structural Biology 2020, 63, 97–105. 10.1016/j.sbi.2020.04.007.

(48) Schnorr, W. E.; Childers, W. S. Biomolecular Condensates: From Bacterial Compartments to Incubator Spaces of Emergent Chemical Systems in Matter-to-Life Transitions. ChemSystemsChem 2024, 6 (4), e202400011. 10.1002/syst.202400011.

(49) Ramšak, M.; Ramirez, D. A.; Hough, L. E.; Shirts, M. R.; Vidmar, S.; Eleršič Filipič, K.; Anderluh, G.; Jerala, R. Programmable de Novo Designed Coiled Coil-Mediated Phase Separation in Mammalian Cells. Nat Commun 2023, 14 (1), 7973. 10.1038/s41467-023-43742-w.

(50) Ghirlanda, G. et al. De Novo Design of a D2-Symmetrical Protein That Reproduces the Diheme Four-Helix Bundle in Cytochrome Bc1. Journal of the American Chemical Society 2004, 126.

(51) Erdős, G.; Pajkos, M.; Dosztányi, Z. IUPred3: Prediction of Protein Disorder Enhanced with Unambiguous Experimental Annotation and Visualization of Evolutionary Conservation. Nucleic Acids Research 2021, 49 (W1), W297–W303. 10.1093/nar/gkab408.

(52) Hoang, Y.; Azaldegui, C. A.; Dow, R. E.; Ghalmi, M.; Biteen, J. S.; Vecchiarelli, A. G. An Experimental Framework to Assess Biomolecular Condensates in Bacteria. Nat Commun 2024, 15 (1), 3222. 10.1038/s41467-024-47330-4.

(53) Cheng, X.; Guinn, E. J.; Buechel, E.; Wong, R.; Sengupta, R.; Shkel, I. A.; Record, M. T. Basis of Protein Stabilization by K Glutamate: Unfavorable Interactions with Carbon, Oxygen Groups. Biophys J 2016, 111 (9), 1854–1865. 10.1016/j.bpj.2016.08.050.

(54) Zimmerman, S. B.; Trach, S. O. Estimation of Macromolecule Concentrations and Excluded Volume Effects for the Cytoplasm of Escherichia Coli. Journal of Molecular Biology 1991, 222 (3), 599–620. 10.1016/0022-2836(91)90499-V.

(55) Stringer, M. A.; Cubuk, J.; Incicco, J. J.; Roy, D.; Hall, K. B.; Stuchell-Brereton, M. D.; Soranno, A. Excluded Volume and Weak Interactions in Crowded Solutions Modulate Conformations and RNA Binding of an Intrinsically Disordered Tail. J. Phys. Chem. B 2023, 127 (26), 5837–5849. 10.1021/acs.jpcb.3c02356.

(56) Zosel, F.; Soranno, A.; Buholzer, K. J.; Nettels, D.; Schuler, B. Depletion Interactions Modulate the Binding between Disordered Proteins in Crowded Environments. Proceedings of the National Academy of Sciences 2020, 117 (24), 13480–13489. 10.1073/pnas.1921617117.

(57) Dębski, D.; Smulik, R.; Zielonka, J.; Michałowski, B.; Jakubowska, M.; Dębowska, K.; Adamus, J.; Marcinek, A.; Kalyanaraman, B.; Sikora, A. Mechanism of Oxidative Conversion of Amplex® Red to Resorufin: Pulse Radiolysis and Enzymatic Studies. Free Radical Biology and Medicine 2016, 95, 323–332. 10.1016/j.freeradbiomed.2016.03.027.

