## Supplementary information for "A Cofactor-Activated Molecular Switch for Condensate Biogenesis and Catalysis *in Escherichia coli*"

***for***

**Cofactor-Driven Phase Separation of a Coiled-Coil Peptide Polymer in *Escherichia coli***

**Materials and Methods**

**Bacterial Strains, Plasmids, and Primers**

All expression constructs were assembled by NEBuilder HiFi DNA Assembly (New England Biolabs, Ipswich, MA, USA) from PCR-amplified fragments and verified by Sanger sequencing (Azenta/GeneWiz). The M1 coding sequence was originally cloned into pET28a(+) between the XhoI and XbaI restriction sites. To generate M1–iLOV, the iLOV sequence was amplified from the donor plasmid pET(His6×-iLOV-TEVp_RGG_T1)^1^ with primers incorporating 3′ overlap with the C-terminus of M1, and the M1 coding region was similarly amplified with overlap primers. The two PCR products were joined by Gibson assembly and re-cloned into pET28a(+). The single-helix control construct M1SH–iLOV was assembled using the same strategy. Because the two helical-domain sequences share high nucleotide identity, the M1SH insert was codon-optimized to minimize secondary structure and recombination during cloning; high GC content in certain regions necessitated inclusion of GC Enhancer and 5% (v/v) DMSO in the relevant PCR amplifications. All final plasmids were transformed into *E. coli* BL21(DE3) (New England Biolabs, Ipswich, MA, USA) by heat-shock chemical transformation and selected on LB agar supplemented with 50 µg mL^−1^ kanamycin. Working stocks were maintained in *E. coli* NEB 5-alpha (New England Biolabs, Ipswich, MA, USA) for plasmid propagation. Plasmid DNA was isolated using a QIAprep Spin Miniprep Kit (QIAGEN N.V., Germantown, MD, USA) and sequence-verified prior to use.

**Protein Expression and Purification**

Expression strains were recovered from glycerol stocks stored at −80 °C and streaked onto LB agar plates containing 50 µg mL^−1^ kanamycin. A single colony was used to inoculate 10 mL LB medium and grown overnight at 37 °C with orbital shaking at 200 rpm. The overnight culture was diluted into 1 L LB medium in a 2 L baffled flask and grown at 37 °C until the optical density at 600 nm (OD_600_) reached 0.6–0.8. Recombinant protein expression was induced by addition of isopropyl β-d-1-thiogalactopyranoside (IPTG) to a final concentration of 1 mM, and cultures were grown for a further 6 h at 37 °C with 200 rpm shaking. Cells were harvested by centrifugation at 5,000 × g for 30 min, and cell pellets were stored at −80 °C until purification.

Frozen cell pellets were thawed on ice and resuspended in 30 mL binding buffer (20 mM sodium phosphate, 300 mM NaCl, 10 mM imidazole, pH 8.0). Cells were lysed by sonication on ice using a 30 s on/30 s off duty cycle for a total processing time of 20 min. Cell debris was removed by centrifugation at 15,000 × g for 40 min at 4 °C. Clarified lysate was loaded onto a 5 mL HisTrap HP column (Cytiva, Marlborough, MA, USA) equilibrated with binding buffer and connected to an ÄKTA Pure chromatography system (Cytiva, Marlborough, MA, USA). Unbound material was removed by washing with 10 column volumes (CV) of wash buffer (20 mM sodium phosphate, 700 mM NaCl, 20 mM imidazole, pH 8.0), and His^6^×-tagged protein was eluted with elution buffer (20 mM sodium phosphate, 300 mM NaCl, 500 mM imidazole, pH 8.0). Eluted fractions corresponding to the main UV absorbance peak were pooled and exchanged into binding buffer by passage through a PD-10 desalting column (Cytiva, Marlborough, MA, USA; Sephadex G-25 resin). The desalted sample was further purified by size-exclusion chromatography (SEC) on a HiLoad 16/600 Superdex 75 pg column (Cytiva, Marlborough, MA, USA) equilibrated in binding buffer without imidazole. M1–iLOV eluted at approximately 61 mL and M1SH–iLOV at approximately 80 mL under these conditions. Sample purity was assessed by SDS-PAGE. Total crude yields were 10–15 mg L^−1^ of culture, with purified yields of 8–12 mg L^−1^ after SEC.

**MALDI-TOF Mass Spectrometry**

Purified protein samples were desalted using C18 ZipTips (MilliporeSigma, Burlington, MA, USA) by sequential wetting, equilibration, sample binding, washing, and elution steps according to the manufacturer’s protocol, exchanging samples into 50:50 (v/v) acetonitrile:water containing 0.1% (v/v) trifluoroacetic acid (TFA). Desalted eluates were mixed 1:1 (v/v) with a saturated solution of α-cyano-4-hydroxycinnamic acid (α-CHCA) matrix and spotted onto a polished stainless steel MTP 384 target plate. Mass spectra were acquired on a Bruker RapifleX MALDI-TOF/TOF instrument (Bruker Corporation, Billerica, MA, USA) operating in linear positive-ion mode. The monoisotopic mass of the singly protonated [M+H]^+^ species was recorded.

**UV–Visible Spectroscopy**

Protein concentrations were determined by UV–visible absorbance spectroscopy using a Cary 50 Bio UV–Visible Spectrophotometer (Agilent Technologies, Santa Clara, CA, USA). The molar extinction coefficient of the iLOV chromophore was ε_450_ = 14,200 M^−1^ cm^−1^. Hemin concentrations were determined using ε_385_ = 58,400 M^−1^ cm^−1^ at the Soret maximum. IHP concentrations were determined using an experimentally derived extinction coefficient of ε_385_ = 48,000 M^−1^ cm^−1^. Metalloporphyrin binding to M1–iLOV was assessed by monitoring the bathochromic shift and intensification of the Soret absorption band upon titration of hemin or IHP into apo-protein solution. Data were analyzed as described previously^2^. Because of the sample turbidity, measurements involving condensates were taken using a 0.1mm path length quartz spectrophotometer cell (Starna Cells Inc., Atascadero, CA, USA).

**Preparation of metalloporphyrin–protein complexes and crowding agents**

Fe(III) protoporphyrin IX chloride (hemin; Sigma-Aldrich) was dissolved in 1 M KOH and diluted into assay buffer (20 mM sodium phosphate, 150 mM NaCl, pH 7.4). Concentration was determined spectrophotometrically (ε = 58,400 M⁻¹ cm⁻¹ at the Soret maximum). For metalloporphyrin loading, hemin was mixed with M1–iLOV at a 1:1 molar ratio and incubated at room temperature for 30 min. Formation of the bis-histidine–ligated complex was confirmed by UV–vis spectroscopy as the appearance of a Soret band at 414 nm. Isohematoporphyrin IX (IHP) chloride was prepared identically; its concentration was determined using ε = 48,000 M⁻¹ cm⁻¹ (experimentally measured). IHP binding was confirmed by UV-vis spectroscopy: a Soret band at 407 nm indicates bis-histidine coordination.

Crowding agents were prepared fresh on the day of use. PEG-8000 was dissolved as a 50% (w/v) stock in ultrapure water and protected from light. Sodium L-glutamate monohydrate was dissolved in ultrapure water to the desired concentration and passed through a 0.22 µm membrane filter.

**Surface passivation of microscope coverslips**

Ibidi μ-Slide glass coverslips (#1.5, 0.17 mm) (Ibidi USA, Fitchburg, WI) were cleaned by soaking in 2 M KOH in isopropanol for ≥30 min, rinsing with ultrapure water, and bath sonication in Liquinox detergent solution for 10 min. Residual detergent was removed by repeated rinsing and sonication in ultrapure water, and coverslips were dried overnight at 70 °C.

PEG-silane functionalization solution (5 mg mL⁻¹ in 96% ethanol, 1% v/v glacial acetic acid) was applied as a 50 µL sandwich between two cleaned coverslips and heated at 70 °C for 30 min to promote covalent silane attachment. Coverslip sandwiches were separated by bath sonication in ultrapure water for 10 min, rinsed sequentially with 20% (v/v) ethanol and ultrapure water (2–3 times each), dried, and stored at room temperature until use.

**Circular Dichroism Spectroscopy**

CD spectra were recorded on a JASCO J-815 spectropolarimeter (JASCO Inc., Easton, MD, USA) equipped with a Peltier thermoelectric temperature controller. Spectra were collected from 200 to 260 nm using a 0.1 nm step size and a scan speed of 50 nm min^−1^ in continuous scanning mode, with a detector integration time of 1 s. Each spectrum represents the average of three accumulated scans. For samples containing sodium glutamate and/or PEG-8000, buffer-matched blank spectra (100 mM sodium glutamate and/or 9% (w/v) PEG-8000 in the appropriate buffer) were subtracted to isolate the protein contribution. Measurements were all taken in a closed demountable 0.1mm path length quartz spectrophotometer cell (Starna Cells Inc., Atascadero, CA, USA).

**Confocal Fluorescence Microscopy**

All confocal imaging was performed on a Nikon A1R HD25 laser-scanning confocal microscope (Nikon Instruments Inc., Melville, NY, USA) equipped with a Plan Apo VC 60× water-immersion objective (NA 1.2, DIC N2). Images were acquired in Galvano scanning mode to maximize spatial resolution. iLOV fluorescence was excited at 445 nm and emission collected at 498.8 nm. DAPI fluorescence was excited at 405 nm and emission collected through a 450/50 nm bandpass filter. Resorufin (Amplex Red oxidation product) fluorescence was excited at 561 nm and emission collected at 595 nm. All in vitro condensates were formed on PEGylated 24x60mm #1 coverslips sandwiched on either side of an 8-well silicone isolator with adhesive (Thermo Fisher Scientific, Carlsbad, CA, USA). In vivo *E. coli* cells (1µL of OD^600^ = 0.2–0.4) were immobilized on 1.0% (w/v) agarose pads prepared in M9 minimal medium (low-melt ultrapure agarose). After heating, agarose was cooled at room temperature inside of silicone isolator wells using a precleaned microscope slide (Thermo Fisher Scientific, Waltham, MA, USA) as a base. Following sample addition to the solidified agarose pad and 5 minute incubation period, an uncoated precleaned 24x60mm #1 coverslip was used to cover the agarose pad and silicone isolator prior to imaging.

**DAPI Staining of the E. coli Nucleoid**

*E. coli* BL21(DE3) harboring pET28a(+)-M1–iLOV or pET28a(+)-M1SH–iLOV were grown in LB medium supplemented with 50 µg mL^−1^ kanamycin at 37 °C with 200 rpm shaking until OD^600^ = 0.6–0.8. Protein expression was induced with 1 mM IPTG, and 1 mL aliquots were collected at the indicated time points. Cells were pelleted by centrifugation (10,000 × g, 5 min, room temperature) and washed three times with phosphate-buffered saline (PBS, pH 7.4). Nucleoid staining was performed by resuspension in PBS containing 10 µg mL^−1^ DAPI (4′,6-diamidino-2-phenylindole) for 10 min at room temperature in the dark. Stained cells were washed three times with M9 salt solution, and diluted prior to immobilization and imaging.

**In Vivo Peroxidase Activity Assay**

*E. coli* BL21(DE3) harboring pET28a(+)-M1–iLOV or pET28a(+)-M1SH–iLOV were grown in M9 minimal medium supplemented with 0.2% (w/v) casamino acids and 50 µg mL^−1^ kanamycin at 37 °C with 200 rpm shaking. At OD^600^ = 0.2, δ-aminolevulinic acid (δ-ALA) and FeCl^3^ were added to final concentrations of 500 µM and 250 µM, respectively, to support heme biosynthesis. Expression was induced with 1 mM IPTG at OD^600^ = 0.5–0.6, and cells were cultured for 5 h at 37 °C with 200 rpm shaking. A 1 mL aliquot was harvested by centrifugation (10,000 × g, 5 min, room temperature) and the cell pellet was washed twice with 1× PBS. Washed cells were resuspended in PBS containing 100 µM Amplex Red and 50 µM hydrogen peroxide and incubated for 15 min at room temperature to allow peroxidase-catalyzed substrate oxidation. Cells were washed once with PBS before dilution, immobilization and imaging.

**In Vitro Peroxidase Activity Assay**

M1–iLOV was pre-incubated with IHP for 30 min prior to condensate formation on the coverslip surface. The molecular crowder, PEG-8000, was added after protein and substrates (Amplex Red) had been equilibrated on the slides. Hydrogen peroxide was added to the condensate-containing sample immediately before imaging. Colocalization of iLOV and resorufin fluorescence signals was verified by acquiring sequential images at 445 nm and 561 nm excitation. Images were processed using FIJI/ImageJ. Fluorescence intensities within condensates were quantified by manually drawing regions of interest (ROIs) around individual droplets; approximately 10 droplets were measured per field of view. Mean fluorescence intensities were exported and analyzed in Excel and Mathematica.

**In Vivo Image Analysis and Cell Thresholding**

Cell segmentation and condensate quantification were performed using FIJI/ImageJ with the MicrobeJ plugin. For each image, the fluorescence channel was thresholded using the Huang method, with manual adjustment of threshold boundaries as needed based on the background intensity of the agarose pad. Cell segmentation was restricted to objects satisfying rod-shaped morphology criteria (area and length parameters consistent with *E. coli*) using the MicrobeJ fit-shape function; the transmitted-light channel was used as an aid for accurate cell boundary identification. Medial fluorescence intensity profiles (width = 10 pixels) for all accepted cells were exported as .mat files and processed in MATLAB using a script adapted from the Obermeyer lab (Liao, 2023) to detect intracellular condensates and calculate the fraction of condensate-containing cells per field of view. Threshold parameters within the script were calibrated to the baseline fluorescence intensity of each image. Data were compiled and plotted using Excel and Mathematica.

**Fluorescence Recovery After Photobleaching (FRAP)**

FRAP experiments were performed on a Nikon A1R HD25 laser-scanning confocal microscope (Nikon Instruments Inc., Melville, NY, USA). Five pre-bleach frames were acquired to establish baseline fluorescence intensity. The region of interest (ROI) was photobleached to near-zero intensity using the 445 nm laser at 100% power for 1–2 s. Post-bleach fluorescence recovery was recorded over the indicated time intervals for three ROIs per field: the bleached area, a non-bleached reference region on a separate cell (for in vivo experiments), and a background region. For in vivo experiments, multipolar *E. coli* cells containing bright condensates were preferentially selected to maximize the available fluorescent pool for recovery. Entire condensates were bleached and recovery of the whole condensate area was monitored. Raw fluorescence intensities were exported to Excel; background intensities were subtracted, and in vivo recovery traces were additionally normalized to the reference region to correct for photobleaching during image acquisition. In vitro droplet traces were not reference-normalized.

Normalized in vivo FRAP traces were fit to a monoexponential recovery equation:

F(t) = B + A·(1 − exp(−k·t))

where F(t) is the normalized fluorescence intensity at time t, B is the residual (non-recovered) fraction, A is the mobile fraction amplitude, and k is the first-order recovery rate constant. The half-time of recovery was calculated as t^1/2^ = ln(2)/k. In vitro FRAP traces were fit to a biexponential model to account for distinct fast and slow diffusional populations:

F(t) = B + A^1^·(1 − exp(−k^1^·t)) + A^2^·(1 − exp(−k^2^·t))

Fast and slow diffusion half-times of recovery were calculated using fast and slow recovery rate constants (k^1^ and k^2^) in place of the first-order recovery rate constant (k). All curve fitting and figure preparation were performed in Mathematica.

Supplementary Figures.

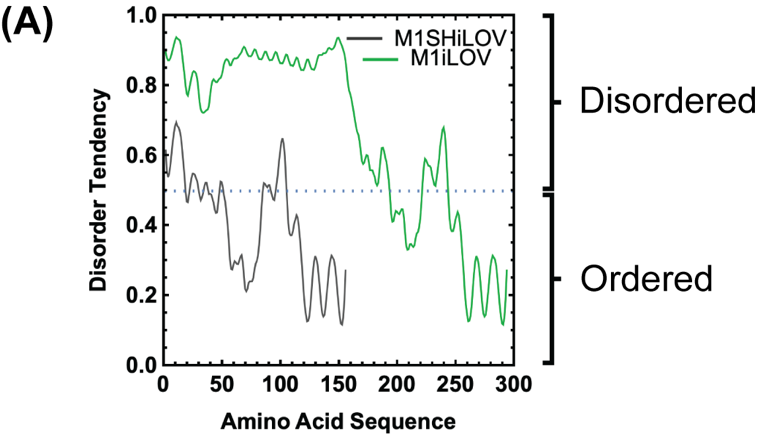

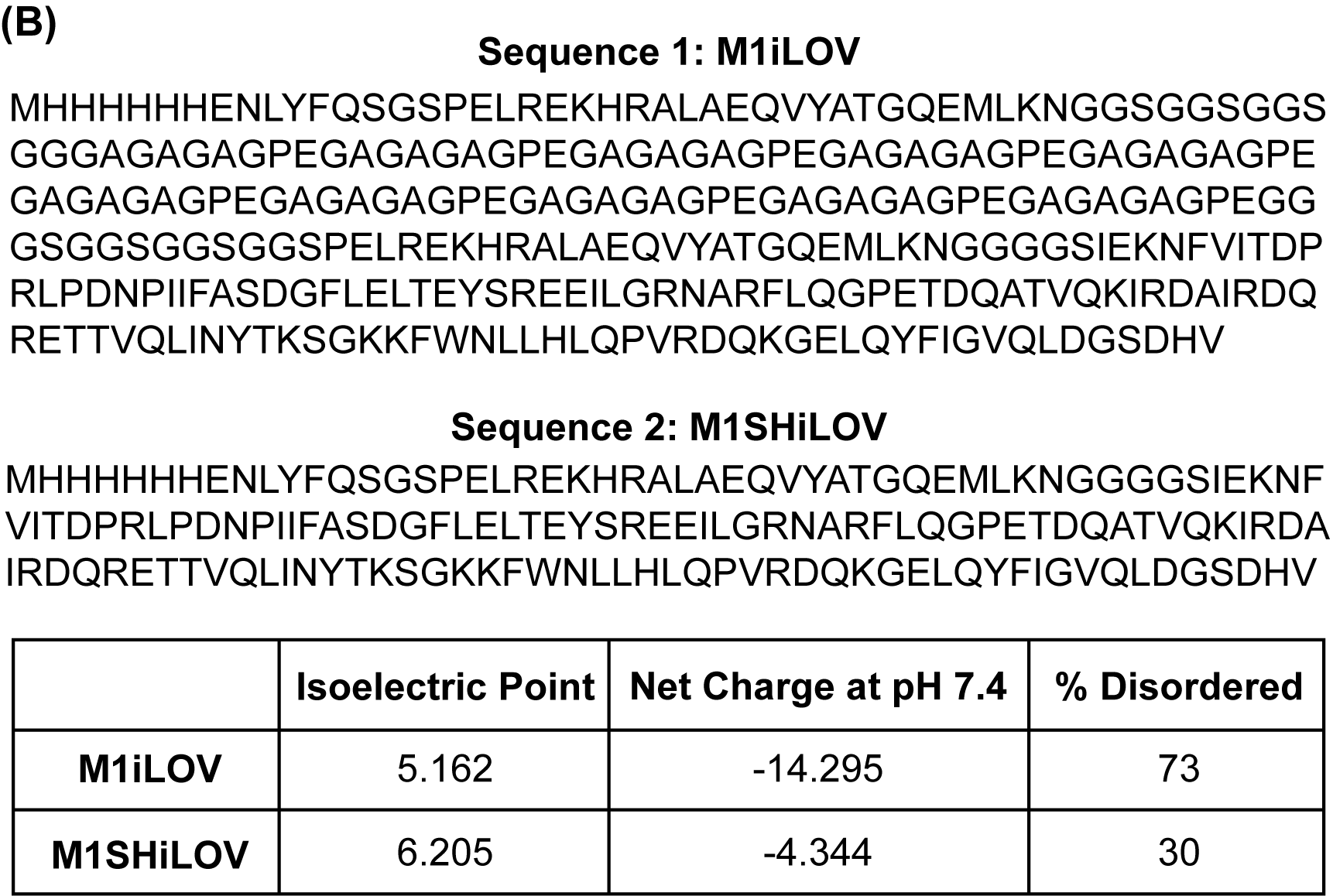

**Figure S1: Primary structure and biophysical analysis of M1iLOV and M1SHiLOV.** (A) IUPred3 analysis of both designed proteins. (B) Primary structure of both uncleaved protein sequences. Biophysical characterization of both sequences was given for contextual understanding of condensate propensity.

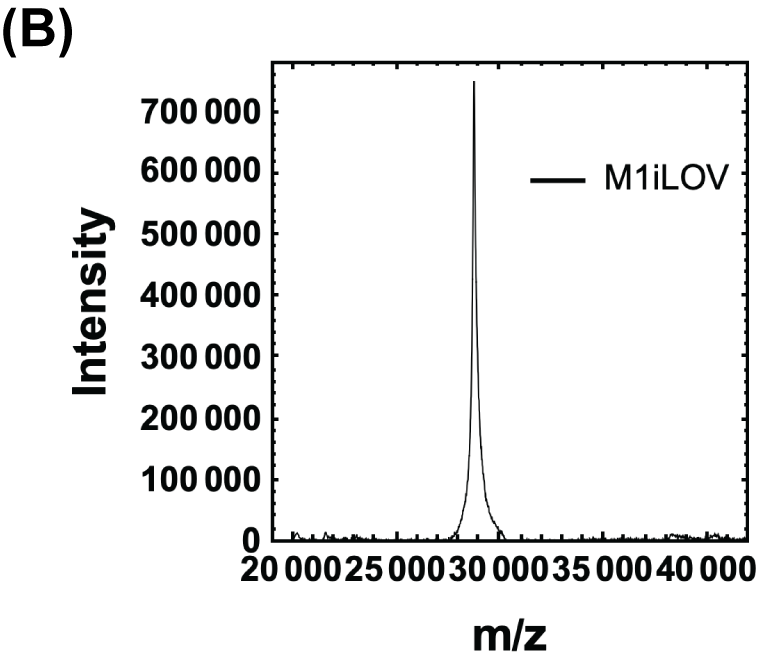

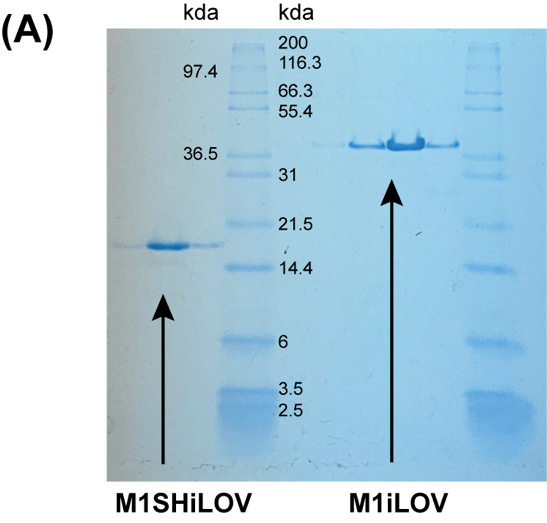

**Figure S2: Purification and characterization of M1iLOV and M1SHiLOV.** (A) A 15% 37.5:1 acrylamide:bisacrylamide polyacrylamide gel showing elution fractions of SEC purified protein (Left of labelled ladder: M1SHilOV; Right of labelled ladder: M1iLOV). Apparent molecular weight is 39.3 kda from gel. (B) Following MALDI mass spectrometery purified M1iLOV was estimated to be around 28.9 kda.

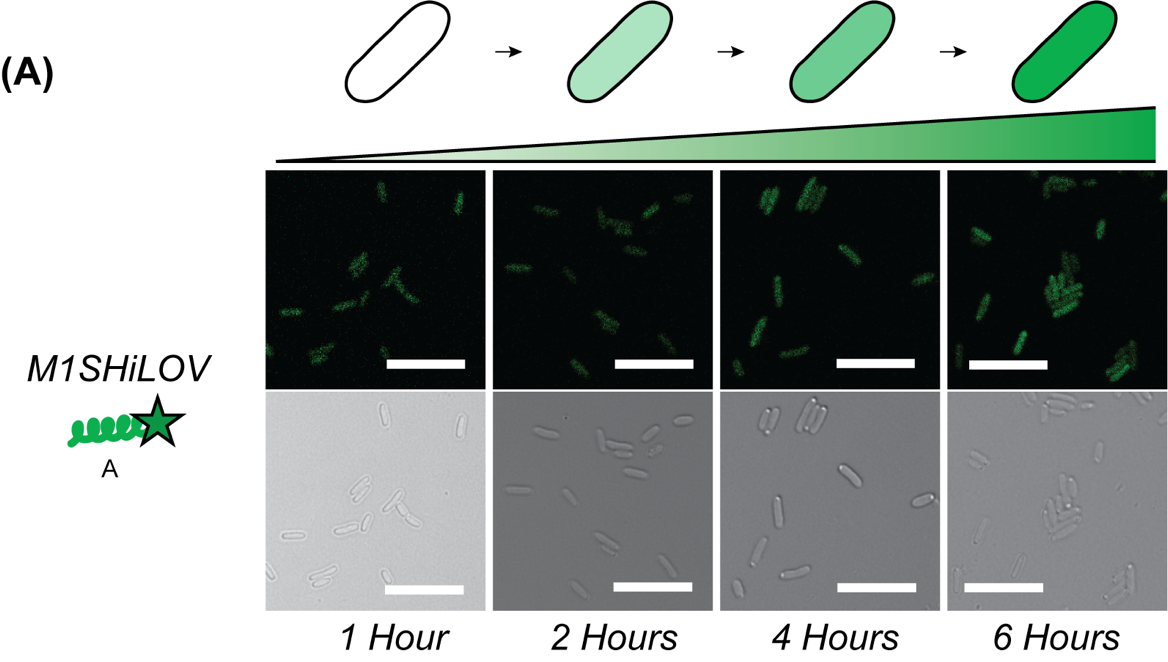

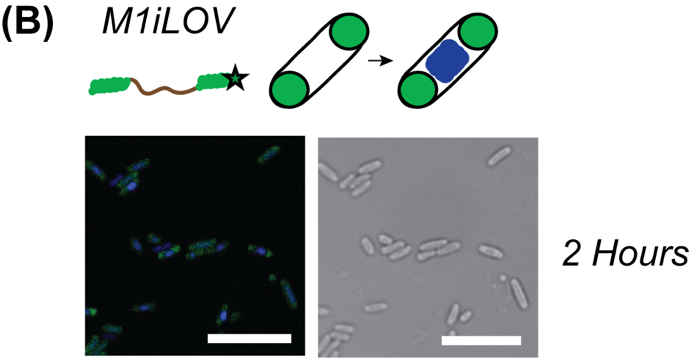

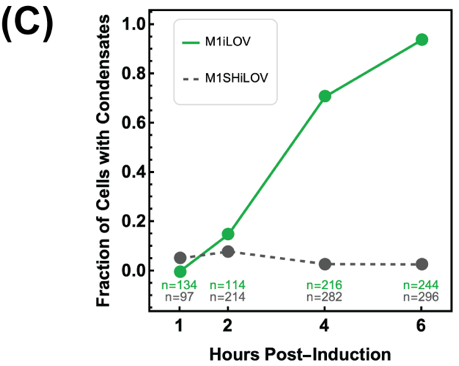

**Figure S3:** (A) Single helix construct (M1SHiLOV) increases concentration in a time-dependent manner without forming biomolecular condensates. Images are shown 1,2,4 and 6 hours after 1mM IPTG induction. Scale bar: 10µm. (B) Triblock construct (M1iLOV) shows commencement of condensate formation with a corresponding compact nucleoid in multiple cells after 2 hours post-induction. (C) Time-dependent comparison of the fraction of cells with condensates between both constructs at 1, 2, 4, and 6 hours post IPTG addition. M1iLOV shows substantial increase in condensate formation between hours 2 and 4.

**Figure S4: M1SH-iLOV doesn’t form foci in E. coli.** Confocal microscopy images showing the absence of biomolecular condensates in the single helix construct (M1SHiLOV) in minimal media across all expression conditions. Images were all taken of E. coli cells collected 5 hours after 1mM IPTG addition at 37 degrees Celsius with shaking. Scale bar: 10µm.

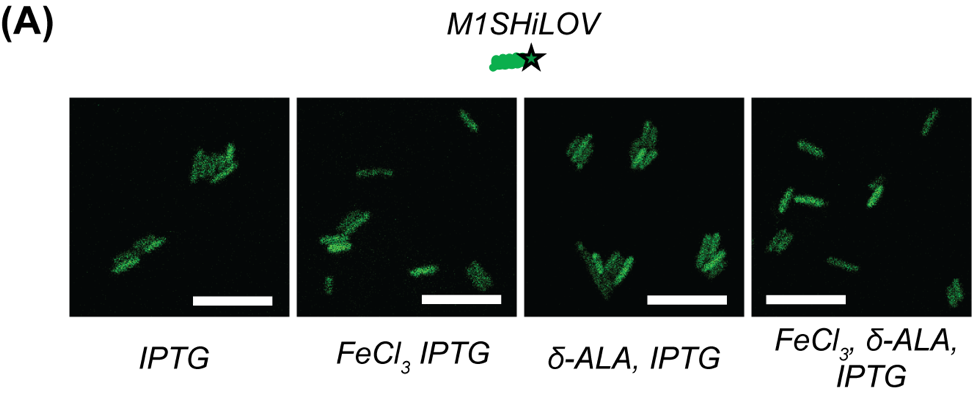

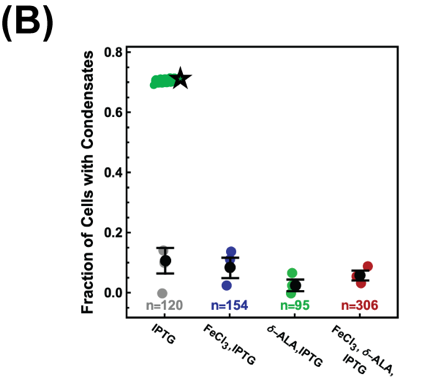

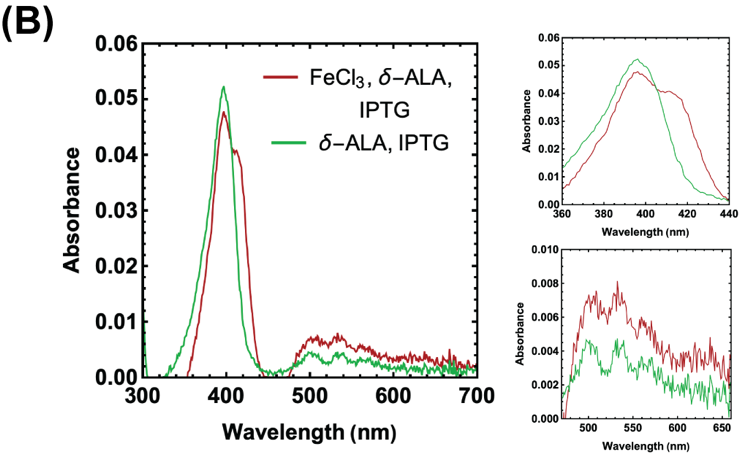

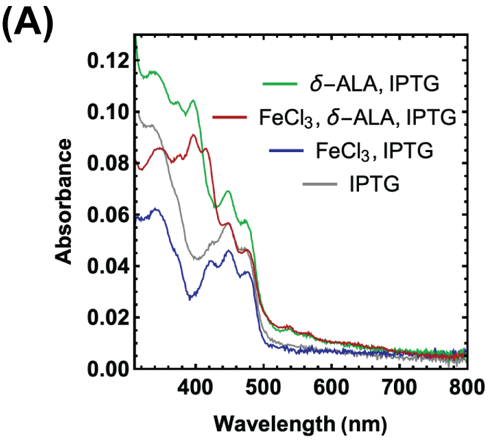

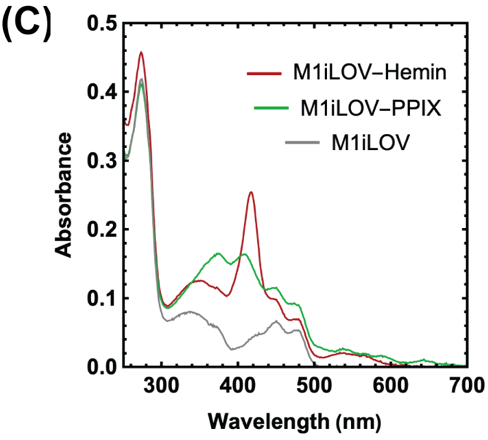

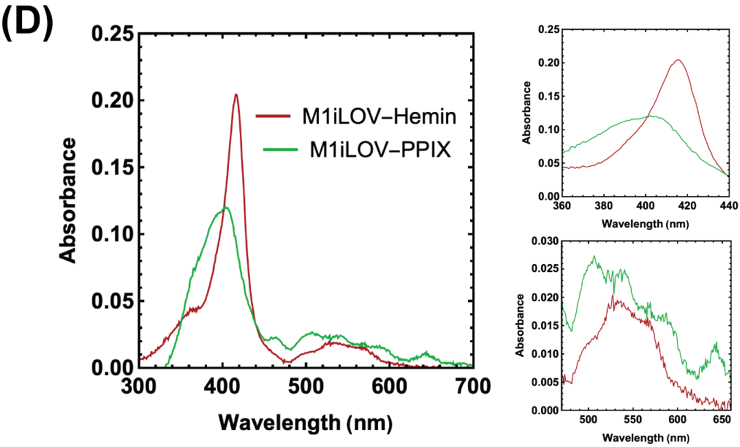

**Figure S5: Heme biosynthesis occurs intracellularly.** (A) UV-visible spectroscopy of the cell lysates of the M1iLOV constructs in various growth conditions after 5 hours of expression. Soret peaks at 395 nm and 414 nm demonstrate protoporphyrin and Fe(III)PPIX formation intracellularly. (B) Spectral subtraction of the two porphyrin forming conditions with the IPTG condition was done to highlight porphyrin formation. Soret and Q-bands were shown on the right. (C) M1iLOV was reconstituted with hemin (red) and PPIX (green) to compare to the lysate spectra. Apo M1iLOV was shown alongside for comparison (gray). (D) Apo M1iLOV spectra was subtracted from the other spectra for a simplified comparison. Soret and Q-bands were shown on the right.

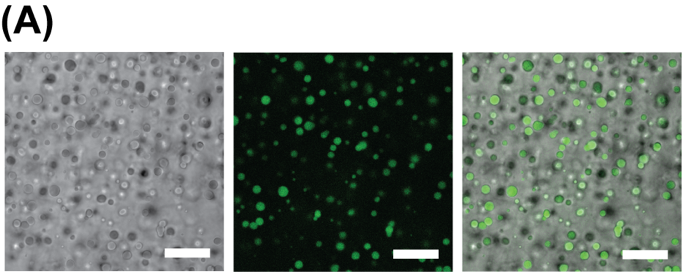

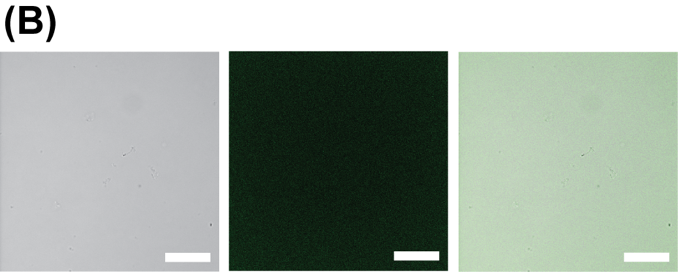

**Figure S6: M1-iLOV and M1SH-iLOV with IHP.** (A) M1-iLOV forms biomolecular condensates in conditions that mimic the intracellular environment with IHP addition; IHP was used to ensure metalloprotein solubility at high concentration. Experimental condition:100µM M1-iLOV- 350µM IHP, 100mM sodium glutamate, 9% PEG-8000 (left); 100µM M1SH-iLOV-350µM IHP, 100mM sodium glutamate, 9% PEG-8000 (right). (B) Similar concentrations of M1SHiLOV do not form biomolecular condensates at similar conditions. Scale bar: 20µm.

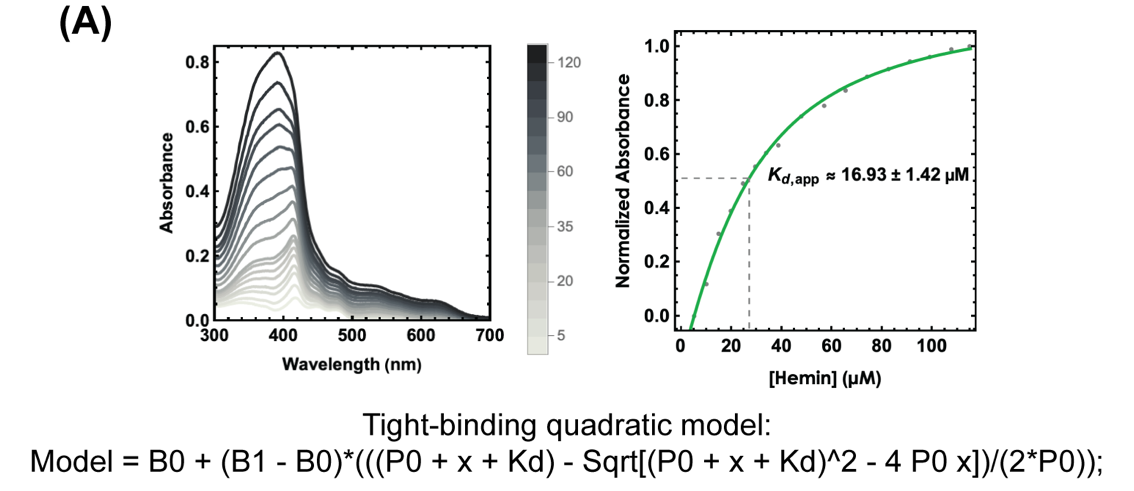

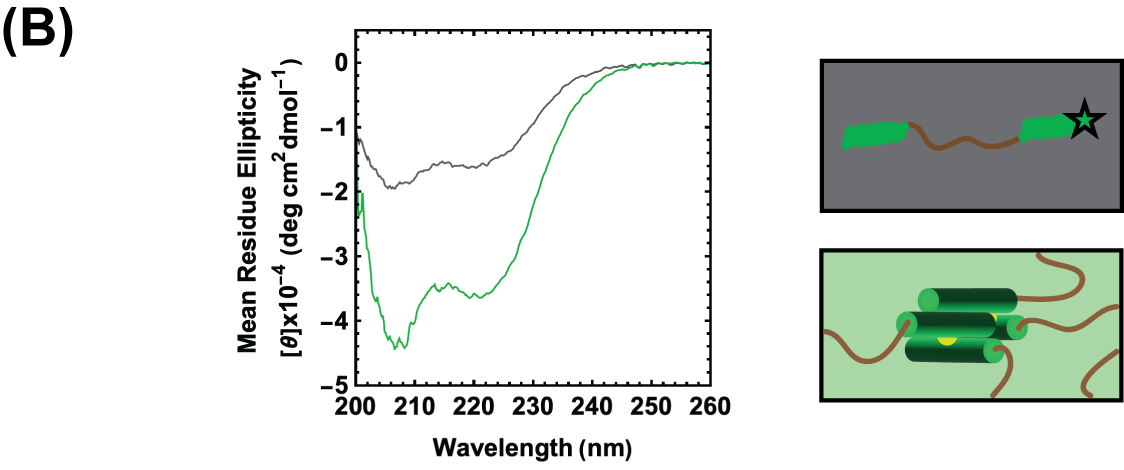

**Figure S7: Hemin-induced folding of M1-iLOV in solution.** (A) Hemin was titrated into 20µM M1iLOV protein at 5µM increments between 0-40µM and at 10µM increments between 40-130µM (left). Absorbance values were adjusted to determine a relationship between bound and unbound states (415nm/390nm*-1) and plotted on the y-axis against hemin concentration (right). A tight-binding quadratic model was used to determine Kd,app. (B) Circular Dichroism of 20µM M1iLOV without hemin (gray) was compared to 20µM M1iLOV with 200µM hemin (green). 9% PEG-8000 and 100mM sodium glutamate was present in both conditions, with the sodium glutamate spectrum subtracted from both spectra.

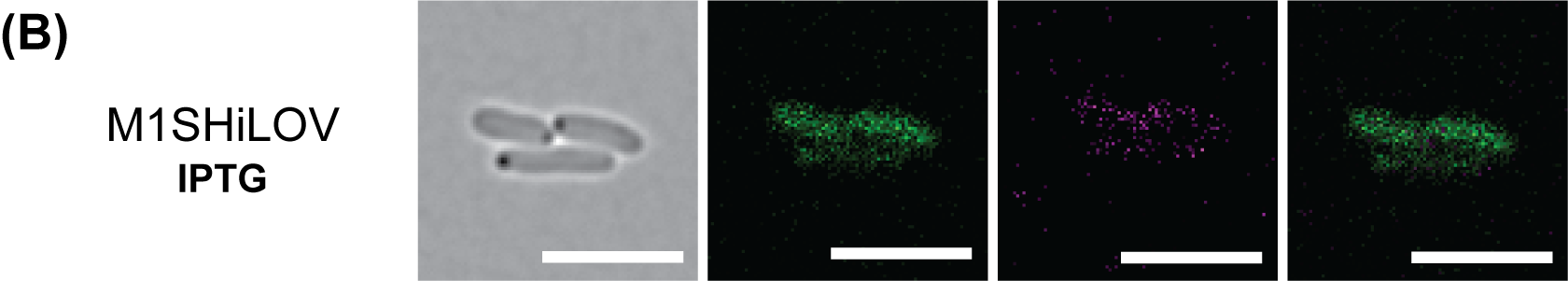

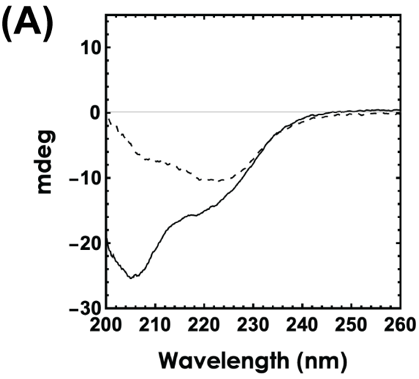

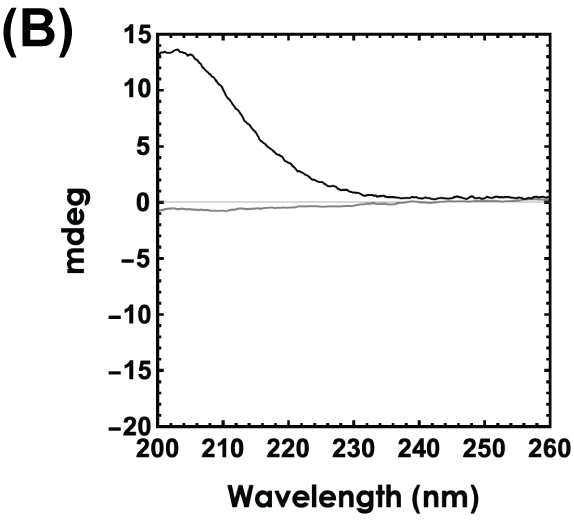

**Figure S8: CD spectra of M1-iLOV in solution.** (A) Millimolar sodium glutamate addition alters secondary structure of the coiled coil alpha helices in 20µM M1iLOV (dotted line) as compared to normal 20µM M1iLOV (solid line). M1iLOV with glutamate has an increased 222nm:208nm ratio. (B) Cd spectra of 9% PEG-8000 and 100mM sodium glutamate used as baseline.

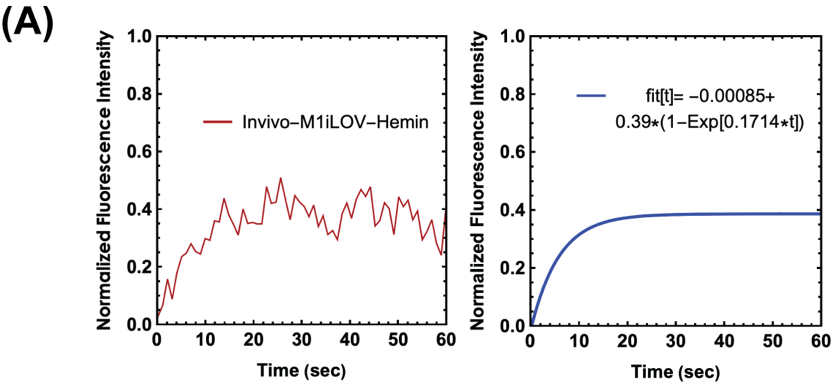

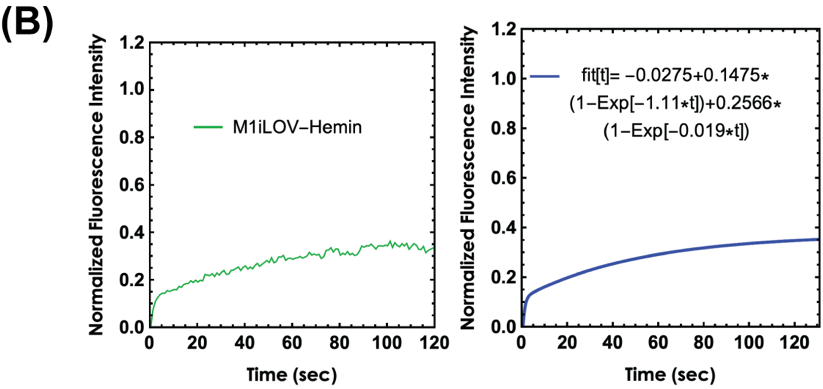

**Figure S9: In vivo and in vitro FRAP exponential fit visualization.** (A) Averaged normalized FRAP intensities for the M1iLOV-heme condensates in vivo were fit to a monoexponential recovery equation to approximate the first order recovery rate constant (k = 0.1714). (B) Averaged normalized FRAP intensities for M1iLOV-hemin in vitro condensates that were partially bleached were fit to a bi-exponential model showing both fast (k1 = 1.11) and slow (k2 = 0.019) second order recovery rate constants.

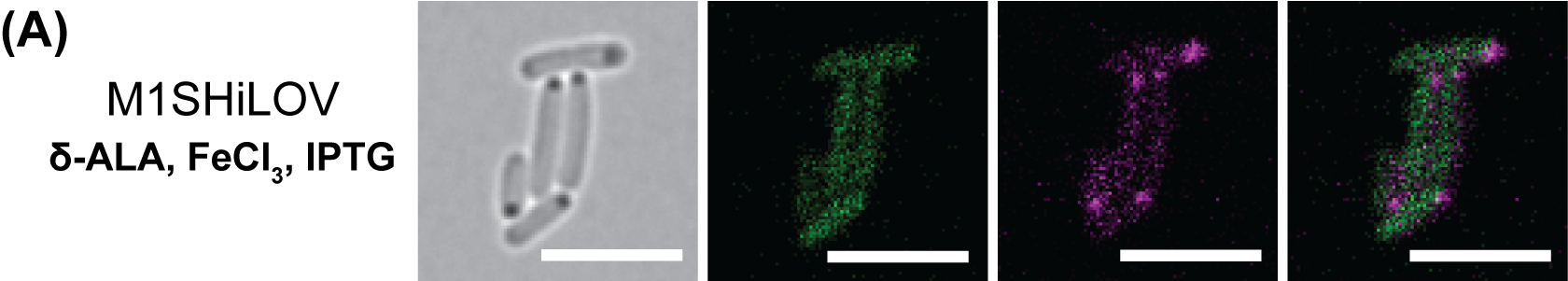

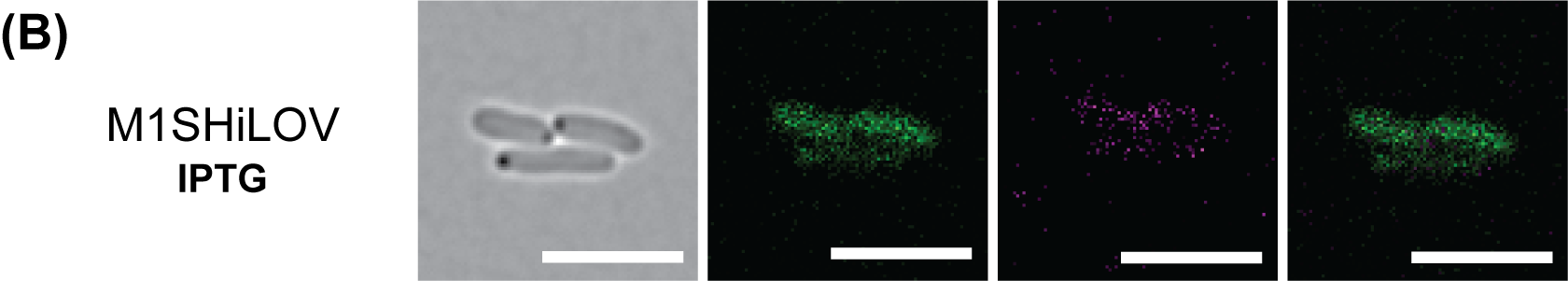

**Figure S10: Peroxidase activity for the single helix construct.** (A) Following (500µM) δ-ALA and (250µM) FeCl3 and 1mM IPTG supplementation, M1SHiLOV was expressed for 5 hours prior to imaging. Foll0wing treatment with 100µM Amplex Red and 50µM hydrogen peroxide, resorufin formation occurred in a unipolar localization pattern. (B) M1SHiLOV was expressed for 5 hours following only 1mM IPTG supplementation and imaged. Resorufin formation appears to be diminished intracellularly without porphyrin. Scale bar: 5µm.

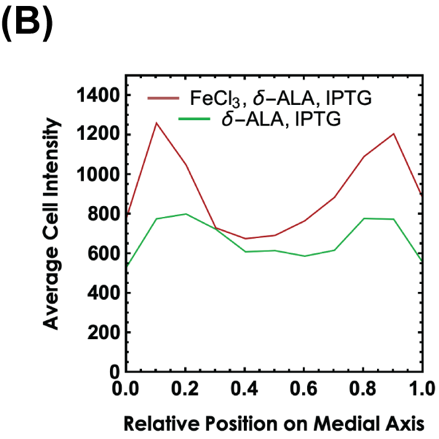

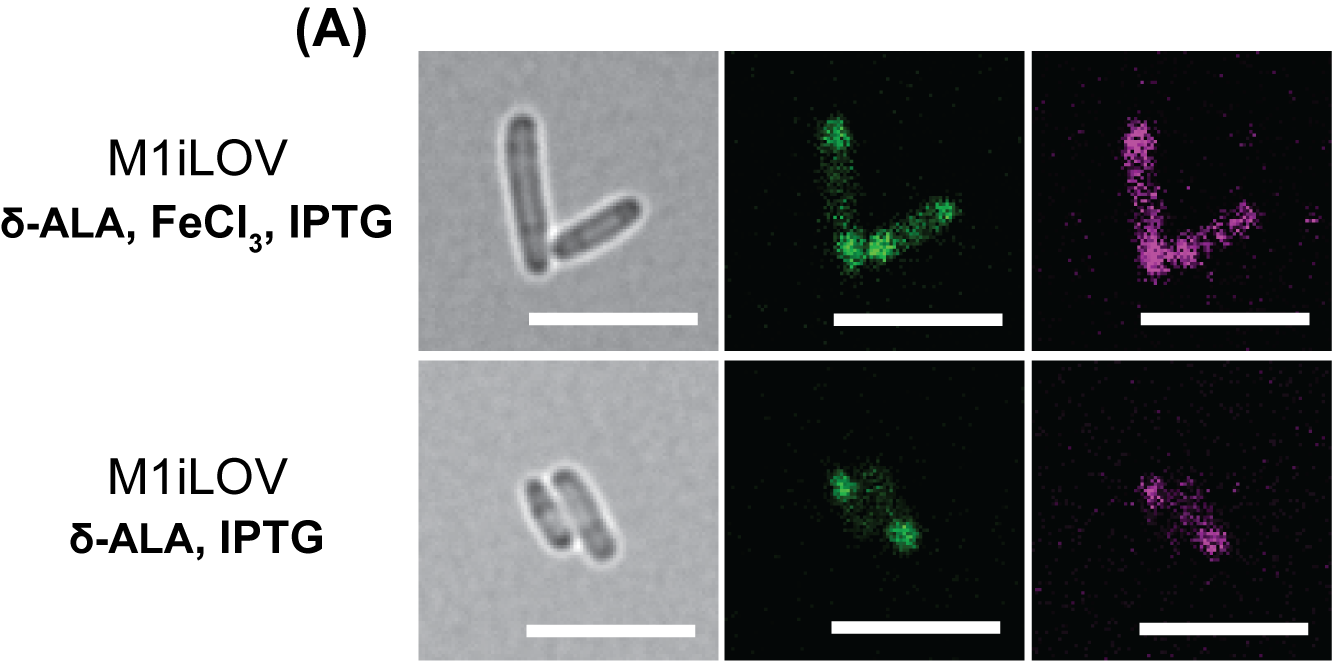

**Figure S11: Background resorufin formation at 0 added hydrogen peroxide.** (A) Confocal microscopy of M1iLOV biomolecular condensates in vivo with and without FeCl3 addition during growth. Only Amplex Red (100µM) was added for incubation to measure endogenous hydrogen peroxide. Scale bar: 5µm (B) Normalized cell intensities on various positions along the E. coli medial axis with (n=35) and without (n=33) FeCl_3_ addition. Intensities for cells with FeCl_3_ addition are larger in the polar regions indicating increased resorufin formation, likely due to increased Fe(III)PPIX.

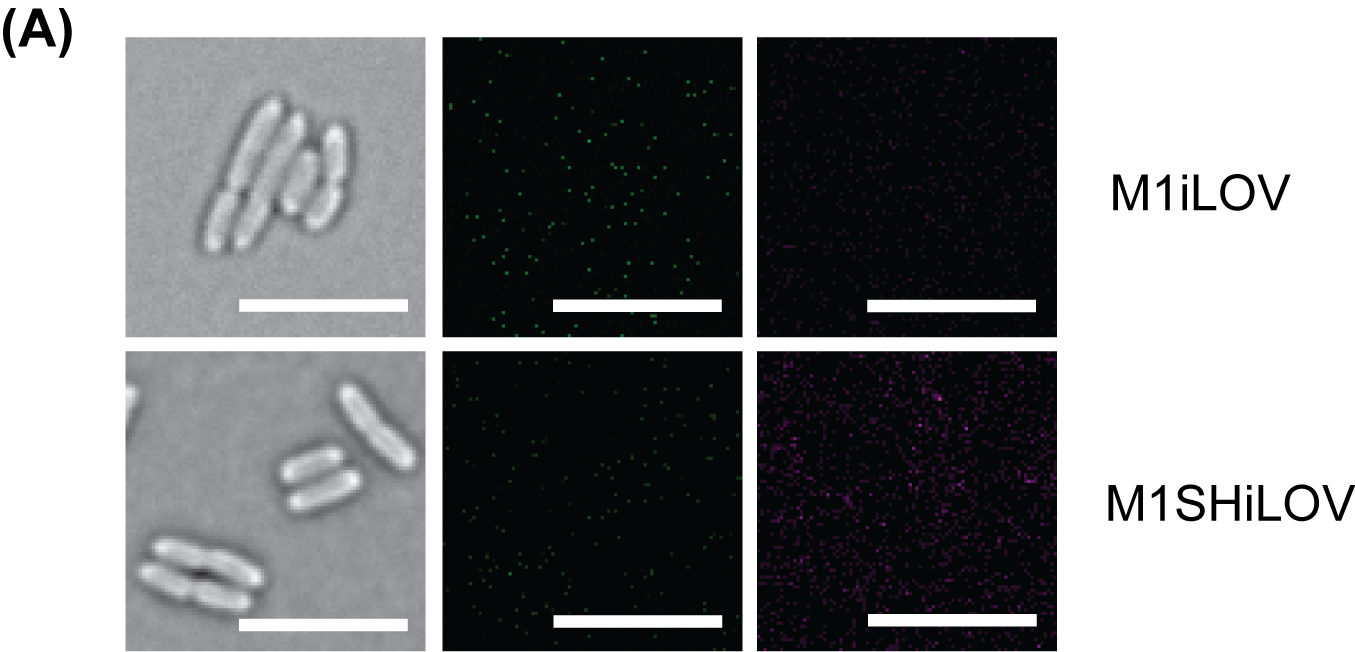

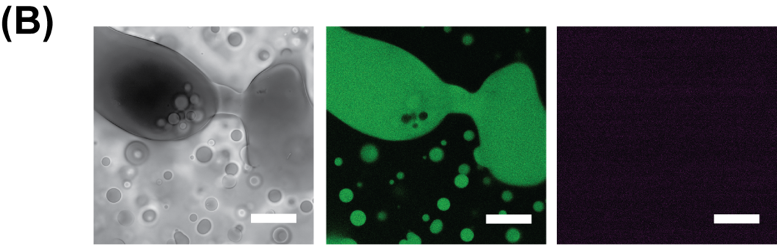

**Figure S12: Peroxidase activity negative controls.** (A) Cells with 0 mM IPTG added during logarithmic growth underwent the same peroxidase activity protocol, yielding negligible 445nm or 561nm fluorescence signal during imaging of M1iLOV (top) and M1SHiLOV (bottom). Scale bar: 5µm. (B) In vitro droplets, without any Amplex Red added, were imaged showing no flavin contribution to the 561nm fluorescent signal. Concentrations were the following: 100µM M1iLOV, 350µM IHP, 9% PEG-8000 (w/v%), 100mM Sodium Glutamate. Scale bar: 20µm

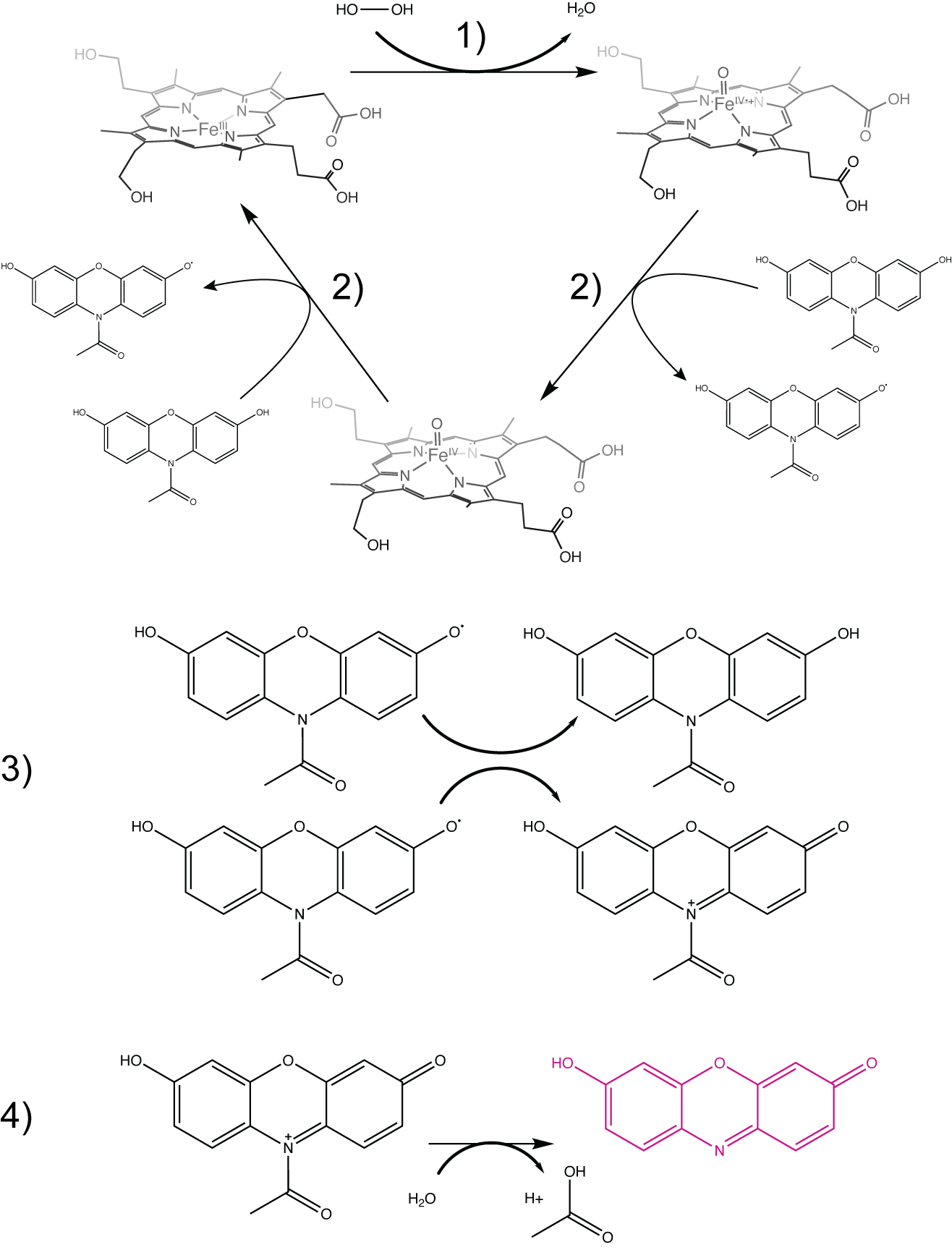

**Scheme S1: Mechanism for heme-mediated radical oxidation of Amplex Red to resorufin.** (1) Heme reacts with hydrogen peroxide to form a high oxidation state intermediate with water as a byproduct. (2) Activated heme takes an electron from a hydroxyl group on Amplex Red leaving a cation radical. (3) Two Amplex Red radicals undergo a disproportionation reaction forming an unoxidized and oxidized Amplex Red molecule. (4) The acetyl group is removed during a secondary hydrolysis step on the oxidized Amplex Red molecule, yielding the fluorescent resorufin product.
